# An empirical model to predict survival curve and relative biological effectiveness after helium and carbon ion irradiation based solely on the cell survival after photon irradiation

**DOI:** 10.1101/2020.06.19.161836

**Authors:** David B. Flint, Scott J. Bright, Conor H. McFadden, Teruaki Konishi, Daisuke Ohsawa, Alisa Kobayashi, Simona F. Shaitelman, Gabriel O. Sawakuchi

## Abstract

**Purpose:** To develop an empirical model to predict radiosensitivity and relative biological effectiveness (RBE) after helium (He) and carbon (C) ion irradiation with or without DNA repair inhibitors.

**Methods:** We characterized survival in eight human cancer cell lines exposed to 6 MV photons and to He- and C-ions with linear energy transfer (LET) values of 2.2-60.5 keV/μm to verify that the radiosensitivity parameters (D_5%_, D_10%_, D_20%_, D_37%_, D_50%_ and SF_2Gy_) correlate linearly between photon and ion radiation with or without DNA-PKcs or ATR inhibitors. Then, we parameterized the LET response of the parameters governing these linear correlations up to LET values of 225 keV/μm using the data in the Particle Irradiation Data Ensemble (PIDE) v3.2 database, creating a model that predicts a cell’s ion radiosensitivity, RBE and ion survival curve for a given LET on the basis of the cell’s photon radiosensitivity. We then trained this model using the PIDE database as a training dataset, and validated it by predicting the radiosensitivity of the cell lines we exposed to He- and C- ions with LET ranging from 2.2-60.5 keV/μm.

**Results:** Radiosensitivity to ions depended linearly with radiosensitivity of photons in the range of investigated LET values and the slopes and intercepts of these linear relationships within the PIDE database vary exponentially and linearly, respectively. Our model predicted ion radiosensitivity (e.g., D_10%_) within 5.1–21.3%, RBE_D10%_ within 5.0-17.1%, and ion mean inactivation dose within 6.7-25.1% for He- and C-ion LET ranging from 2.2-60.5 keV/μm.

**Conclusions:** Radiosensitivity to He- and C-ions depend linearly with radiosensitivity to photons and can be used to predict ion radiosensitivity, RBE and cell survival curves for clinically relevant LET values from 2.2–60.5 keV/μm, with or without drug treatment.

**SUMMARY:** We present a new empirical model capable of predicting clonogenic cell survival of cell lines exposed to helium and carbon ion beams. Our model is based on an observed linear correlation between radiosensitivity to ions and photons across a wide range of LET values. This linear correlation can be used to predict ion RBE, radiosensitivity, and the cell survival curve for a given LET all based on a cell’s photon survival curve.

## INTRODUCTION

Cancer therapy using carbon (C) ion beams has a number of benefits compared to conventional photon beams, including inherently superior depth-dose distributions (1,2), less dependence on tumor oxygenation status (3), and increased biological effectiveness (1,2). But, while much progress has been made understanding the physical properties of ions, much remains unclear about how ion response depends on cell biology and to what extent the physical and biological mechanisms governing radiosensitivity to ions interact with one another in determining ion radiosensitivity.

Several models have been proposed to explain the widely-characterized variations in relative biological effectiveness (RBE) on radiation quality, including the microdosimetric kinetic model (MKM) (4,5), which explains the increasing RBE values for increasing linear energy transfer (LET) values in terms of microdosimetric changes, and the local effect model (LEM) (6–8), which explains the variations in RBE in terms of how increasing the LET affects the heterogeneity of the microscopic dose distribution. However, these models do not incorporate much biological information, despite numerous observations that biological factors including tumor histology (9,10), genotype (9,10), cell cycle phase (10–13), and DNA damage repair capacity (10,13,14) greatly affect cellular radiosensitivity, and that the LET response varies greatly between cell lines (15). Furthermore, while these models are used to predict the RBE in clinical settings (10), they have not been tested to predict the ion response of cell lines treated with radiosensitizing drugs, which is critical given recent advances in combining novel DNA repair inhibitor or chemotherapy drugs with radiation.

Thus, the extent to which our current ion RBE models can accurately predict the response of cells with differing genotypes, histologic subtypes, DNA damage repair capacities, or subjected to drug treatments is unknown. But, in agreement with Suzuki et al. (16), our data show that independently of how these biological factors might govern a cell’s inherent (photon) radiosensitivity, the relationship between a cell’s radiosensitivity to ions, including helium (He-) and C-ions, and its radiosensitivity to photons is linear. And thus, if this linear relationship is known for a particular ion LET, a cell’s radiosensitivity to that ion LET, and thus its RBE, can be predicted based only on a cell’s photon radiosensitivity. This premise forms the basis of the model we present here, and therefore our work represents a simple, easily implementable empirical model that predicts ion radiosensitivity and RBE while implicitly accounting for biological variability and the effects of drugs within its framework due to their modulation of the model’s input: cell radiosensitivity to photons.

## METHODS AND MATERIALS

We selected eight human cancer cell lines of different histologic subtypes, genotypes, and capacity for DNA repair (H460, H1299, BxPC-3, PANC-1, AsPC-1, Panc 10.05, M059K and M059J) to quantify cellular survival after radiation. These cell lines span wide range of radiosensitivities from D_10%,photon_ = (1.37±0.02) Gy for the radiosensitive M059J (glioblastoma) cell line, to D_10%,photon_ = (8.56±0.19) Gy for the radioresistant Panc 10.05 (pancreatic cancer) cell line. The M059J cell line, which is vastly more radiosensitive than the M059K (glioblastoma) cell line established from the same tumor (17), is deficient in the protein DNA-PKcs, which renders them deficient in DNA double strand break (DSB) repair via non-homologous end joining (NHEJ) (18), while the H460 (lung cancer) cell line has wild type TP53 (19) in contrast to the other cell lines which are TP53 mutants. Further details of these cell lines can be found in Supplemental Method 1. All cell lines for this work were obtained from the American Type Culture Collection (ATCC) and independently authenticated at the University of Texas MD Anderson Cancer Center. The cells were cultured in incubators at 37°C and 5% CO_2_.

We exposed the cell lines to He- and C-ions with dose-weighted LET values of 2.2, 7.0, and 14.0 keV/μm (He-ions) and 13.5, 27.9 and 60.5 keV/μm (C-ions) at the Heavy-Ion Medical Accelerator in Chiba (HIMAC) (Chiba, Japan), and to 6 MV photons at The University of Texas MD Anderson Cancer Center (Houston, Texas, USA) (Supplemental Method 3). We performed clonogenic assays to quantify survival for each cell line at each LET (Supplemental Method 4) to characterize their radiosensitivity. H460, H1299, PANC-1 and Panc 10.05 cell lines were also treated the DNA repair inhibitors NU7441, which inhibits DNA-PKcs, which is essential in NHEJ repair, and AZD6738 which inhibits the protein ATR, which is essential in DNA damage response. H1299 was additionally treated with B02 which inhibits Rad51, which is essential in homologous recombination (HR) repair (Supplemental Method 5).

We used the dose required to achieve surviving fractions of 5%, 10%, 20%, 37%, and 50%, as well as the survival fraction at 2 Gy derived from the cell survival curves to quantify cell radiosensitivity after photon or ion irradiations and refer to these parameters as R_photon_ = [D_5%,photon_, D_10%,photon_, D_20%,photon_, D_37%,photon_, D_50%,photon_ and SF_2Gy,photon_] and R_ion_ = [D_5%,ion_, D_10%,ion_, D_20%,ion_, D_37%,ion_, D_50%,ion_ and SF_2Gy,ion_], respectively. All clonogenic cell survival data collected in this work were done in duplicate or triplicate and repeated at least two times, except the data collected for the cell lines treated with inhibitors, for which only a single He- or C-ions experiment was performed in duplicate. However, because for a given cell line treated with inhibitors, three cohorts were used (e.g. DNA-PKcsi, ATRi, DMSO vehicle), the location of the trends derived from this dataset have similar statistical power to three independent experiments performed with each untreated cell line. And so while this dataset has greater variance than the cell lines not treated with inhibitors, because it contains so many individual data points, we can use it to confirm whether the trends in the drug-treated cell lines agree with the untreated cell lines.

Statistical analyses were performed in MATLAB 2017 (Mathworks, Natick, MA) and Graph Pad Prism 7 (Graph Pad, San Diego, CA). Error bars represent the standard error propagated from the fitted survival curve parameters (α and β), including their covariance, to the radiosensitivity parameters estimated from them. The confidence intervals represent the uncertainty from our fit parameters, including their covariance, propagated into our predictive function, calculating the 95% confidence intervals as ±1.96 times the standard error of the prediction.

## RESULTS

### Ion radiosensitivity parameters correlate with photon radiosensitivity parameters

Suzuki et al. (16) noted that D_10%,ion_ is linearly correlated with D_10%,photon_ for C-ions with LET values of 13.3 and 77 keV/μm. Our survival data for cell lines exposed to He- and C-ions (LET ranging from 2.2−60.5 keV/μm) show that this linear relation holds not only for D_10%_ but also for D_5%_, D_20%_, D_37%_, D_50%_ and SF_2Gy_ (Fig. 1, Supplemental Results 1) with R^2^ values ranging from 0.722-0.999 (Supplemental Results 2).

**Fig. 1.**
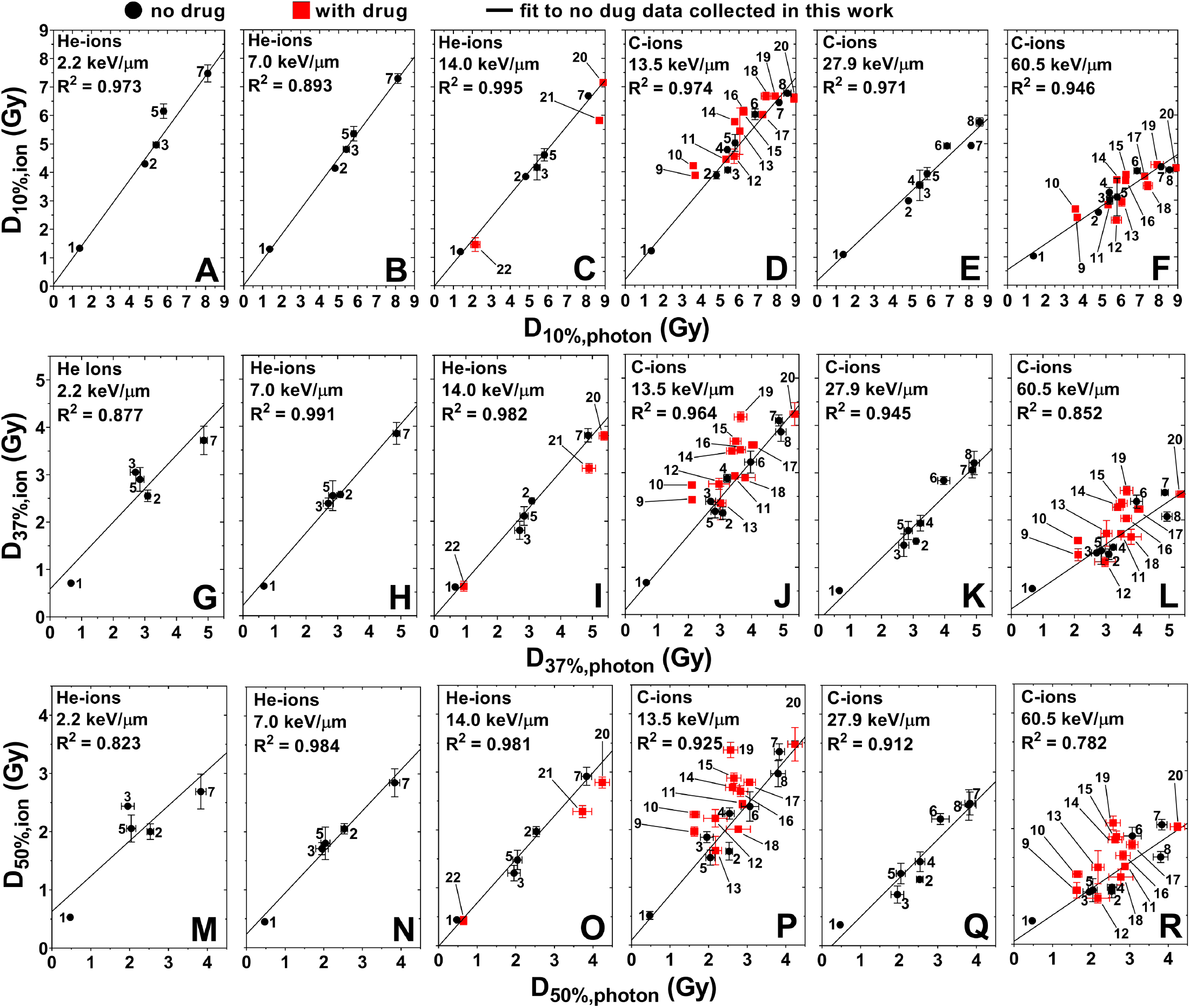
(A-F) D_10%,ion_ vs. D_10%,photon_, (G-L) D_37%,ion_ vs. D_37%,photon_, and (M-R) D_50%,ion_ vs. D_50%,photon_ for He- and C-ions in order of increasing LET from 2.2 to 60.5 keV/μm. Black circles and red squares represent data of cell lines treated with radiation alone and drug in combination of radiation. Lines represent linear fits to the data of cell lines exposed to radiation alone. The results for D_5%_, D_20%_ and SF_2Gy_ are given in Supplemental Results 2. 1: M059J, 2: H460, 3: M059K, 4: AsPC-1, 5: BxPC-3, 6: PANC-1, 7: H1299, 8: Panc 10.05, 9: H460+ATRi (0.1 μM), 10: H460+DNA-PKcsi (0.1 μM), 11: H460+DMSO, 12: Panc 10.05+ATRi (0.1 μM), 13: Panc10.05+DNA-PKcsi (0.1 μM), 14: PANC-1+DNA-PKcsi (0.1 μM), 15: PANC-1+DMSO, 16: PANC-1+ATRi (0.1 μM), 17: H1299+DNA-PKcsi (0.1 μM), 18: Panc 10.05+DMSO, 19: H1299+ATRi (0.1 μM), 20: H1299+DMSO, 21: H1299+Rad51i, and 22:H1299+DNA-PKcsi (1 μM).

### The linear relationship between ion and photon radiosensitivity varies regularly with LET

Although Suzuki et al. (16) previously observed a linear relationship between ions and photons, they did not comment on how the slopes and intercepts of these correlations depend on LET. It is apparent from our data that the slopes decrease with increasing LET.

To confirm the extent to which this holds for all LET values, we analyzed the survival data in the Particle Irradiation Data Ensemble (PIDE) version 3.2 database (20) for cells exposed to He- and C-ions. We binned these data into LET bins of width 2-10 keV/μm, over LET values ranging from 1.8-125 keV/μm for He- ions and 12.16-225 keV/μm for C-ions (Supplemental Method 6). We then plotted R_ion_ vs. R_photon_ for each LET bin, fit the data with a linear function and extracted the slopes and intercepts. We noted that the slopes and intercepts of R_ion_ vs. R_photon_ vary regularly with LET up to at least 125 keV/μm for He-ions and 225 keV/μm for C-ions (Fig. 2). A one phase exponential decay (R^2^ = 0.8981 for D_10%_) and a linear function (R^2^ = 0.4578 for D_10%_) provide good fits for the slope and intercept versus LET, respectively. These functions are empirical and do not represent any intended theoretical description.

**Fig. 2.**
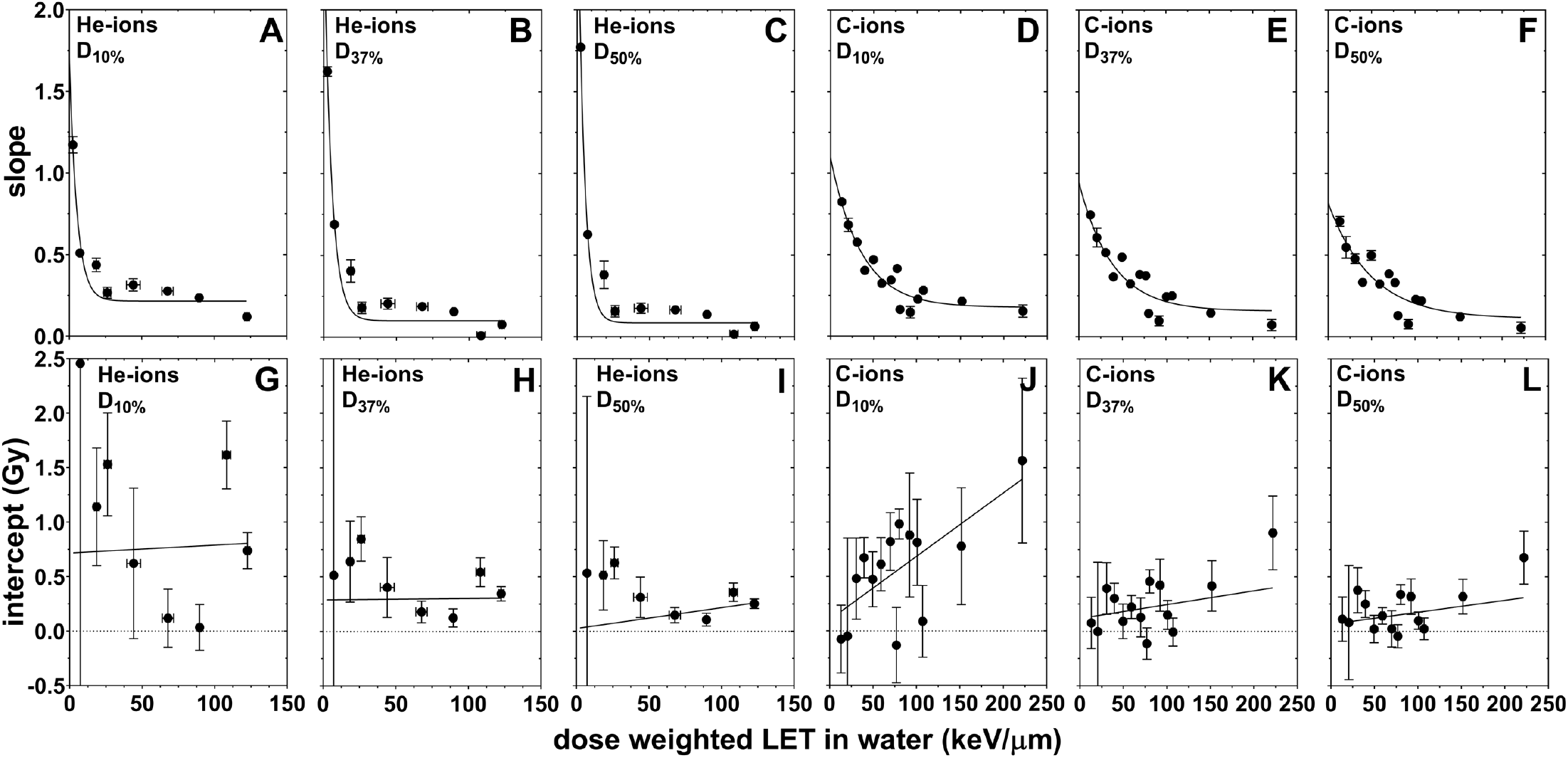
We used the PIDE database (20) to determine the linear trends for several parameters of the survival curve (shown here D_10%_, D_37%_ and D_50%_) for several cell lines (n=42 for He-ions and n=91 for C-ions) and LET values (2.2-225 keV/μm). The linear functions for each LET can be described in terms of the slope and intercept, which in turn can be plotted as functions of LET (A-L). The results for D_5%_, D_20%_ and SF_2Gy_ are given in Supplemental Results 3.

Our formalism can thus be generalized into the following function of 5 parameters, c, d, f, g and h that predict R_ion_ from R_photon_ for a given LET:

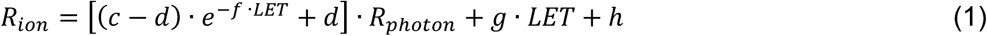

This function scales a cell’s radiosensitivity to photons linearly for a fixed ion LET to give its radiosensitivity to ions, allowing the slope and intercept of this linear scaling to vary exponentially and linearly with LET, respectively. With this form, the data from the PIDE database can be fit without the need for binning to determine the free parameters, and thus we fit the response of all the cell lines in the database to this function for LET values up to 201 keVμm for He-ions (R^2^=0.7657 for D_10%_) and 225 keV/μm for C-ions (R^2^=0.8435 for D_10%_). The values of these parameters are given in Supplemental Results 4.

### Prediction of the linear relationship between ion and photon radiosensitivity

Using our model, we predicted the relationship between ion and photon radiosensitivity for 2.2, 7.0 and 14.0 keV/μm for He-ions and 13.5, 27.9 keV/μm and 60.5 keV/μm for C-ions, and compared these predictions to the measured response of H460, H1299, BxPC-3, PANC-1, AsPC-1 Panc 10.05, M059K and M059J cell lines (Fig 3). These LET values and cell lines were not used to determine the parameters of our model and therefore served as a validation cohort. Our measured data agreed within a root mean square percentage error (RMSPE) of 9.6%, 5.1%, 8.9%, 7.3%, 6.0%, 21.3% at 2.2, 7.0, 14.0 (He-ions), 13.5, 27.9 and 60.5 keV/μm (C-ions) for D_10%_, respectively. The RMSPE for D_5%_, D_20%_, D_37%_, D_50%_ and SF_2Gy_ are shown in Supplemental Results 5.

**Fig. 3.**
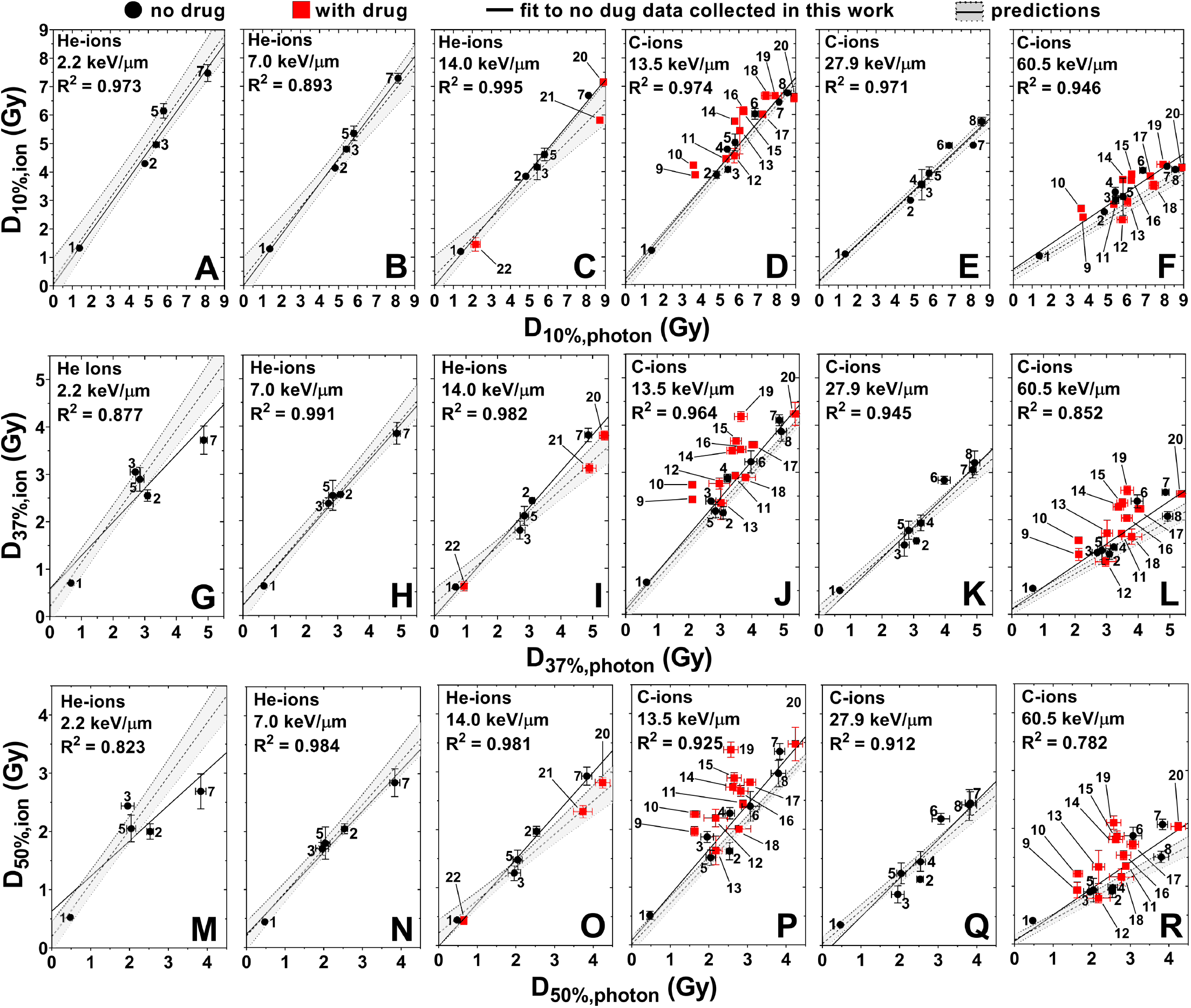
Prediction (with 95% confidence interval) and measured data for the linear relation between D_10%_ (A-F), D_37%_ (G-L) and D_50%_ (M-R) for ions vs. photons. Numbers indicate the cell line and are given in the caption of Fig. 1. Results for D_5%_, D_20%_ and SF_2Gy_ are given in Supplemental Results 5.

### Prediction of ion RBE

RBE is defined as the ratio of the photon dose to the ion dose required to achieve the same biological endpoint:

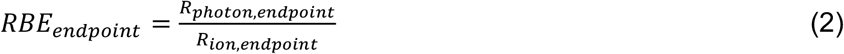

And so, we can substitute our expression for R_ion_ into the expression for RBE to yield the predictive function:

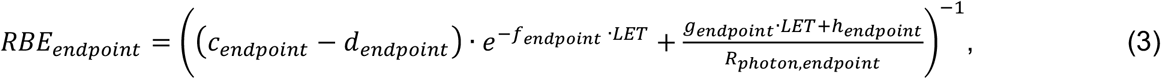

which gives the RBE in terms of our model parameters and R_photon_. Our measured RBE values agreed with this function’s predictions within a RMSPE of 5.0%, 5.3%, 8.1%, 6.8%, 5.7%, 17.1% at 2.2, 7.0, 14.0, 13.5, 27.9 and 60.5 keV/μm for RBE_D10%_, respectively (see Supplemental Results 6 for other parameters and plots of RBE_D5%_ and RBE_D20%_) (Fig. 4).

**Fig. 4.**
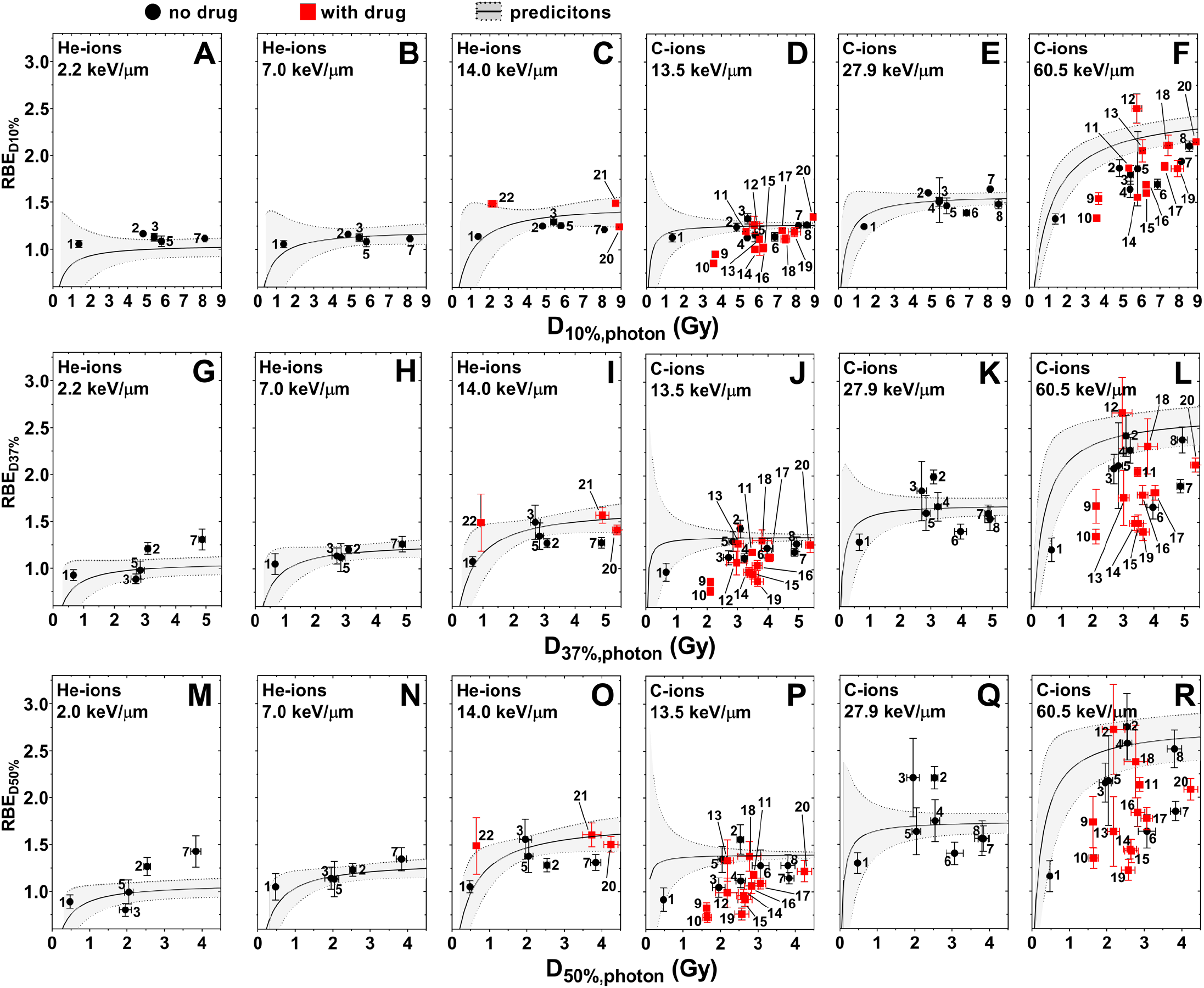
Predicted (with 95% confidence interval) and measured RBE as functions of D_10%,photon_ (A-F), D_37%,photon_ (G-L) and D_50%,photon_ (M-R). Numbers indicate the cell line and are given in the caption of Fig. 1. These cell lines and LET values were not used to determine the linear functions and served as a validation of the model. D_5%_, D_20%_ and SF_2Gy_ are in Supplemental Results 6.

### Prediction of ion survival curves

Using the survival curve for photons (the α_photon_ and β_photon_ values from the linear quadratic model), it is straightforward to calculate all the elements of R_photon_. R_ion_ can then easily be predicted for a given LET using our model. Thus, as our model predicts 6 points on the ion survival curve (D_5%,ion_, D_10%,ion_, D_20%,ion_, D_37%,ion_, D_50%,ion_ and SF_2Gy,ion_), its predictions can be fit to the linear quadratic model (or in principle other models) to predict the ion survival curve (Fig. 5). To quantify the agreement between our predicted and measured survival curves, we integrated the curves to calculate the mean inactivation dose 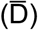. Our model predicts 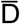 within a RMSPE of 13.6%, 6.7%, 11.8%, 12.6%, 10.1% and 25.1% at 2.2, 7.0, 14.0 (He-ions), 13.5, 27.9 and 60.5 keV/μm (C-ions), respectively.

**Fig. 5.**
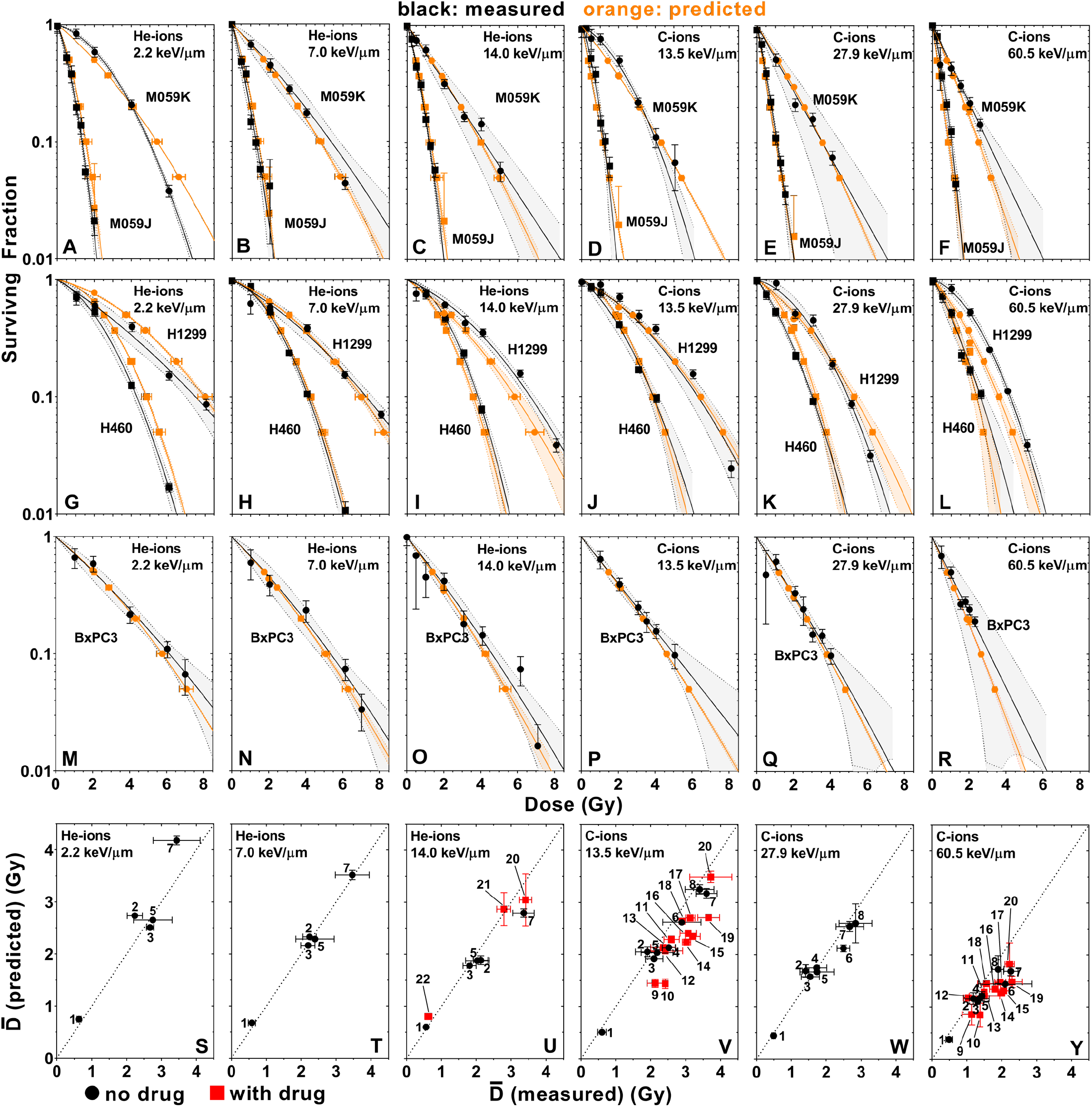
Predicted (orange, with 95% confidence interval) and measured (black, with 95% confidence interval) survival curve for M059K and M059J cells (A-F), H460 and H1299 cells (G-L), and BxPC-3 cells (M-R) exposed to He- and C-ions. Shaded areas are 95% confidence intervals. (S-Y) Predicted versus measured mean inactivation doses 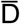 of survival curves from A-R, as well as for the other cell lines curves that are provided in Supplemental Results 7. Numbers indicate the cell line and are given in the caption of Fig. 1.

### Prediction of cell lines treated with ion and DNA repair inhibitors

We compared the response of the DNA repair inhibor treated cell lines to ion verus photon exposure, which we superimposed upon (Fig. 1-Fig. 5) to demonstrate the trends in the data. We do not report R^2^ for these data since each point corresponds to a single independent experiment, and so most of the variance in these data can be attributed to experimental uncertainy. Nevertheless, the agreement of the trends in these data with the trends observed in the untreated cell lines (Fig. 6) indicates that it is possible to predict the response of cell lines treated with DNA repair inhibitors using our model.

**Fig. 6:**
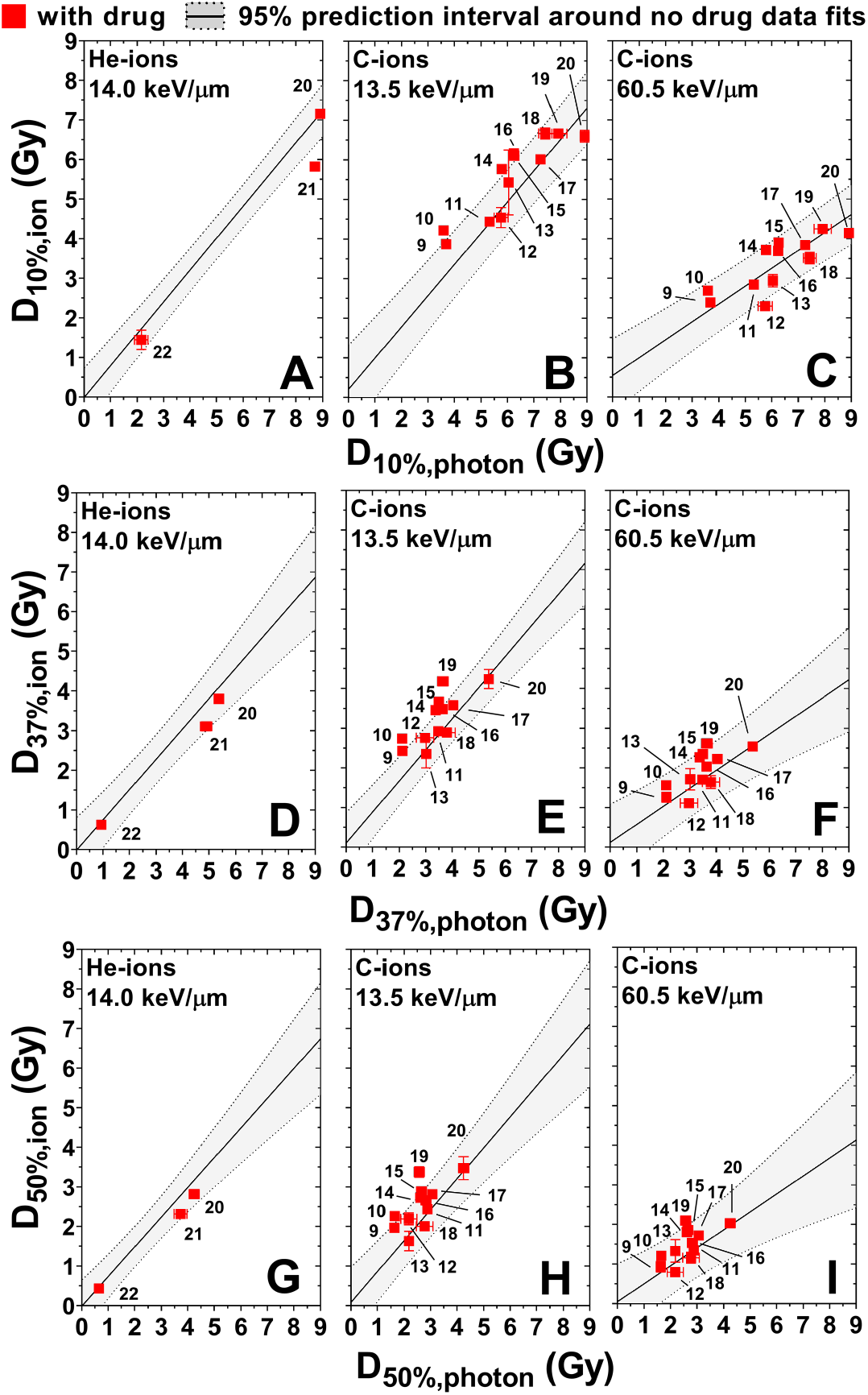
Measured ion versus photon radiosensitivity for cells treated with DNA repair inhibitors (red) superimposed on the 95% prediction interval derived from the data not treated with drugs (gray) for the parameters D_10%_ (A-C), D_37%_ (D-F) and D_50%_ (G-I). Numbers indicate the cell line and are given in the caption of Fig. 1. Results for D_5%_, D_20%_ and SF_2Gy_ are given in Supplemental Results 8.

## DISCUSSION

Ion LET is understood to modulate radiosensitivity through differing patterns of energy deposition: higher LET radiations are more densely ionizing, which results in an increased DNA-double strand break (DSB) yield and clustered DSB yield per unit dose (21) and, consequently, increased RBE. But the LET effect depends also on ion species. For the same LET, different ions will have differing DNA damage yields due to differing track ionization densities, e.g., a He-ion will have a higher DSB and clustered DSB yield compared to a C-ion of the same LET due to its much denser track (21). This generally leads to higher RBE values at the same LET for an ion of lower charge (21). One approach to reconcile the response of different ions is to use an ion-independent substitute for beam quality such as the Q factor proposed by Luhr et al (22). However, as we could only validate our model for two ion species (He- and C-ions), and there were sufficient data in the PIDE database to model each ion’s LET response independently, we decided to simply use dose-weighted LET to parameterize the beam quality, fitting the model parameters for each ion separately to account for the effects of different ions.

The strong linear correlations we observed between ion and photon radiosensitivity suggest that a simple proportionality relationship governs an ion’s RBE for a given dose-weighted LET. This is despite the great biological differences between the cell lines in our panel known to modulate radiosensitivity—particularly with respect to histological type and genotype (9,10), DNA repair capacity (10,13,14), anatomical site, tumorigenicity, species of origin–in addition to the pharmacologic inhibition of a number of DNA repair proteins. This suggests that regardless of whatever biological factors govern a cell’s intrinsic radiosensitivity, the physical component of a cell’s radiosensitivity to ions that is dictated by the beam quality can be described by a simple proportionality constant at each dose-weighted LET.

We can understand this proportionality when we consider how the induction of lethal DNA lesions relates to, for example, the quantity D_10%_. Under the framework of the Curtis Lethal-Potentially Lethal (LPL) model (23), a given dose of radiation will induce a number of DNA lesions, some of which are lethal and result in cell death, and some of which are only potentially lethal if not repaired. But D_10%_ occurs well beyond the repair shoulder (where the repair of potentially lethal lesions is most relevant), and so cell death in the vicinity of D_10%_ is dominated by the induction of irreparable, lethal lesions. Consequently, D_10%_ can be understood as the dose resulting in a 90% probability of inducing at least one lethal lesion. Because the number of lesions induced by a given radiation dose follows a Poisson distribution (4), D_10%_ occurs when the number of lethal lesions per cell, η, has an expectation value of ~2.3. But if we assume, as in the LPL model, that in the vicinity D_10%_ the expected number of lethal lesions induced is proportional to dose (23), then D_10%,ion_, and D_10%,photon_ are both proportional to the same quantity, η≈2.3, and therefore they are both proportional to one another. Similar arguments can be made for the other radiosensitivity parameters examined.

As for how this proportionality varies with LET, the number of DSBs per unit dose increases with ion LET over the range of LET values we investigated (21). Thus, ions with higher LET values will produce more DSBs and thus more lethal DNA lesions for a given dose. Accordingly, we see that as the LET is increased, the slope of R_ion_ versus R_photon_ decreases, because less ion dose is required to induce the same number of lethal lesions.

The trend of increasing positive intercepts of R_ion_ versus R_photon_ with LET can be understood when considering the limit where, for instance, D_10%,photon_ approaches zero. In this limit, cells are extremely radiosensitive to the homogenous radiation doses deposited by photons. But to deliver such small doses with ions requires a fluence so low that many nuclei will not be hit by ions. Consequently, the cells whose nuclei are indeed hit will receive a nuclear dose much larger than D_10%,photon_ and almost certainly die, whereas the cells that are not hit will receive no dose, and, disregarding any bystander effects, survive. Therefore, D_10%,ion_ in this limit is simply the dose that results in 10% of nuclei receiving at least one hit. This number must be positive because it is a physical dose and must increase with LET, because as the LET is increased a greater dose is required to achieve the same fluence.

For very low LET values, the intercepts of the R_ion_ versus R_photon_ trends tend towards small, possibly null values, although their uncertainties are too large to assert how they trend with great confidence. However, for a null intercept, RBE would be independent of R_photon_, and tend to a value of (slope of R_ion_ versus R_photon_)^−1^. This prediction, that at very low LET values, RBE becomes increasingly independent of radiosensitivity and tends to a constant value, is in line with the observation by Britten and Murray (24) that RBE is constant for low LET values, even among cells with different DNA repair proficiencies.

Generally, our model predicts a non-linear relationship between ion RBE and photon radiosensitivity, which follows from the linear relationship between radiosensitivity to ions and photons. This relationship predicts higher RBE values for radioresistant cells, lower RBE values for radiosensitive cells, and sub-unity RBE values for extremely radiosensitive cells (Fig. 4). These predictions are consistent with observations of near (and sometimes sub-) unity RBE for radiosensitive cells deficient in DSB repair (8,25–27) and comparatively large RBE values for radioresistant cells (12,28,29).

This nonlinear relationship also predicts RBE values much less than one in the limit where R_photon_ approaches zero. This unintuitive result follows from the fact that for very radiosensitive cells, the dose required to achieve, for example, 10% survival is substantially higher for ions than for photons. This is because in this limit, only very small photon doses, which are approximately homogenous, are required to achieve 10% survival, whereas for ions, a dose large enough to result in at least 90% of the cells being hit at least once is still required, and thus the RBE values are significantly smaller than one. But in the extremely radiosensitive limit, in which the homogeneity of photon doses breaks down, D_10%_ is achieved when the secondary electron fluence produced by the photon beam is equal to the ion fluence required to achieve 90% hits. The RBE in this case would tend to the ratio of the LET values of the photon beam’s secondary electron spectrum to the ion’s, which typically has a value much less than one.

The near unity of RBE values at low LET values also implies that any small LET effect at low LET values is dominated by the biological factors governing a cell’s inherent radiosensitivity. Only when the LET is increased does the RBE gain a dependence on photon radiosensitivity, which suggests that both physical and biological factors contribute to radiosensitivity for high LET ions. Thus, we posit that in the absence of changes in other physical parameters affecting radiosensitivity such as oxic status, ion radiosensitivity might best be understood by a two-step approach. First, biological factors such as genotype determine a cell’s intrinsic radiosensitivity to low-LET radiation such as photons. Second, a linear relationship scales the cell’s ion sensitivity proportionally to its photon radiosensitivity. This linear scaling, being applied equally to all cell lines, must be biology-independent, and thus represents a purely physical relationship between photon and ion radiosensitivity.

Our work therefore suggests a nuanced relationship between the biological factors governing a cell’s intrinsic radiosensitivity and the physical factors governing the LET effect in determining a cell’s ion radiosensitivity: biological factors determine a cell’s intrinsic radiosensitivity; physical factors determine generally how cell radiosensitivity varies with LET; but biological factors, in determining a particular cell’s intrinsic radiosensitivity, dictate how that cell’s radiosensitivity will vary with ion LET.

A great strength of our model is that it can easily predict the ion survival curve, and thus any parameter that can be derived from a cell survival curve, e.g. α/β, SF_XGy_, D_X%_, 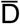, etc. Although this is technically feasible with LEM (6–8) the computations are not straightforward and are too computationally expensive for a clinical setting (30,31). In the case of the MKM model, by contrast, β_ion_ is assumed to be constant (4,5), which differs from experimental observations, and thus only α_ion_ can be predicted. Meanwhile, the simplicity of our model allows the full ion survival curves to be calculated in a single spreadsheet.

Further, the LEM and MKM models are limited in that they require sophisticated Monte Carlo calculations or physical measurements to compute microdosimetric quantities such as the saturation corrected dose-mean specific energy of the beam (in the case of MKM) (4,5) or the local dose distribution within the cell nucleus (in the case of LEM). By contrast, our model requires only the photon survival curve (also required for LEM and MKM) and dose-weighted LET, which is much easier to compute than the physical quantities required by the LEM and MKM models. Notably, the data in the PIDE database contains both monoenergetic beams and mixed fields but reports the dose-weighted LET values, and thus this parameter alone may be sufficient to characterize the beam in the context of our model.

Unique to our model is that it can predict the response of cell lines irradiated with ions in combination with DNA repair inhibitors including DNA-PKcs, ATR and Rad51 inhibitors. This is an important aspect of our model because of ongoing clinical trials involving DNA-repair inhibitors combined with radiation [DNA-PKcs (e.g., NCT02516813) (32), ATM (e.g., NCT03423628) (33), ATR (e.g., NCT02223923, NCT04052555, NCT02567422, and NCT03641547) (34), PARP (e.g., NCT02229656, NCT03212742, NCT03109080, and NCT01589419)]. Because our model implicitly incorporates the biological factors governing cells’ intrinsic radiation response through their radiosensitivity to photons, we hypothesize that it might also predict the response of cells exposed in combination with other DNA repair inhibitors or even chemotherapeutic drugs. Our model may also be able to predict the response of hypoxic cells, however, since hypoxia changes the relationship between radiation dose and cell death to a different extent for x-rays and ions (3), it may be that the trends for hypoxic cells must be modeled separately from normoxic cells. But in all these areas, further investigation is needed to confirm the applicability of our model.

A significant limitation of our model is that it requires photon survival data as its input. While this does indirectly incorporate biological factors governing a cell’s inherent radiosensitivity into the model’s predictions, in a clinical setting, this type of survival data is rarely available. But at the same time, this limitation is faced when using LEM or MKM, and given that the linear relationship between R_ion_ and R_photon_ is observed in cell lines of various histologic subtypes, anatomical sites, DNA repair capacities, and genotypes, the universality of this phenomenon implies that this is likely the only input we need to characterize a cell line’s biological response.

A final limitation to this work is that although our model was trained over a large range of LET values (up to 201 and 225 keV/μm for He- and C-ions, respectively), the LET range over which we validated our model’s predictions (up to 14 and 60.5 keV/μm for He- and C-ions, respectively) was limited. Very high LET values for a given ion can only be achieved at the end of the ion range, where very high dose and LET gradients render survival measurements difficult to make accurately. We choose to limit our highest LET values to minimize these dosimetric uncertainties arising from the experimental setup, thus ensuring the robustness of our validation dataset. So while it is worth noting the maximum LET values we validated are approximately equal to the dose-weighted LET values in the middle of clinically relevant spread-out Bragg peaks (35), for the portion of the beam at the very end of the range where the LET values are extremely high, further validation needs to the done

## CONCLUSIONS

Our data show that radiosensitivity to ions is linearly related with radiosensitivity to photons and we developed a model based on this linear relation that predicts RBE and the survival curve of several cell lines exposed to He- and C-ions based on their response to photons. We showed that this model can be used to predict the He- and C-ions response in cell lines for which the photon response is known, including in cells exposed in combination with DNA-repair inhibitors of DNA-PKcs, ATR and Rad51, which is unique to our model. Our model also predicts several trends observed in the literature including: (i) the RBE of radiosensitive cells varies little with LET, (ii) the RBE of radioresistant cells varies greatly with LET, (iii) RBE is constant and near unity across cell lines for low LET values, and (iv) there is much greater RBE variation between cell lines at higher LET values. These data suggest that biological factors in addition to physical factors have important roles in determining how cell radiosensitivity varies with LET.

## ACKNOWLEDGEMENTS

This work was supported in part by funds from the Cancer Prevention and Research Institute of Texas grant RP170040 (G.O.S.); the Division of Radiation Oncology, The University of Texas MD Anderson Cancer Center (G.O.S.); the University Cancer Foundation via the Institutional Research Grant program (G.O.S.) and via the Sister Institution Network Fund at The University of Texas MD Anderson Cancer Center (G.O.S.); and the Cancer Center Support (Core) Grant CA016672 to The University of Texas MD Anderson. The authors thank and Christine F. Wogan of the Division of Radiation Oncology at MD Anderson for editing the manuscript.

## REFERENCES

1. Tsujii H, Kamada T, Baba M, et al. Clinical advantages of carbon-ion radiotherapy. New J Phys 2008;10.

2. Kamada T, Tsujii H, Blakely EA, et al. Carbon ion radiotherapy in japan: An assessment of 20 years of clinical experience. Lancet Oncol 2015;16:E93–E100.

3. Nakano T, Suzuki Y, Ohno T, et al. Carbon beam therapy overcomes the radiation resistance of uterine cervical cancer originating from hypoxia. Clin Cancer Res 2006;12:2185–90.

4. Hawkins RB. A statistical theory of cell killing by radiation of varying linear energy transfer. Radiat Res 1994;140:366–74.

5. Hawkins RB. A microdosimetric-kinetic model for the effect of non-poisson distribution of lethal lesions on the variation of rbe with let. Radiat Res 2003;160:61–9.

6. Scholz M, Kraft G. Calculation of heavy-ion inactivation probabilities based on track structure, x-ray-sensitivity and target size. Radiat Prot Dosim 1994;52:29–33.

7. Scholz M, Kellerer AM, Kraft-Weyrather W, et al. Computation of cell survival in heavy ion beams for therapy. The model and its approximation. Radiat Environ Biophys 1997;36:59–66.

8. Weyrather WK, Ritter S, Scholz M, et al. Rbe for carbon track-segment irradiation in cell lines of differing repair capacity. Int J Radiat Biol 1999;75:1357–64.

9. Williams JR, Zhang Y, Zhou H, et al. A quantitative overview of radiosensitivity of human tumor cells across histological type and tp53 status. Int J Radiat Biol 2008;84:253–64.

10. Karger CP, Peshke P. Rbe and related modeling in carbon-ion therapy. Phys Med Biol 2017.

11. Blakely E, Chang P, Lommel L, et al. Cell-cycle radiation response: Role of intracellular factors. Adv Space Res 1989;9:177–86.

12. Tsuboi K, Tsuchida Y, Nose T, et al. Cytotoxic effect of accelerated carbon beams on glioblastoma cell lines with p53 mutation: Clonogenic survival and cell-cycle analysis. Int J Radiat Biol 1998;74:71–9.

13. Okayasu R. Repair of DNA damage induced by accelerated heavy ions--a mini review. Int J Cancer 2012;130:991–1000.

14. Genet SC, Maeda J, Fujisawa H, et al. Comparison of cellular lethality in DNA repair-proficient or -deficient cell lines resulting from exposure to 70 mev/n protons or 290 mev/n carbon ions. Oncol Rep 2012;28:1591–6.

15. Blakely EA, Ngo FQH, Curtis SB, et al. Heavy-ion radiobiology – cellular studies. Adv Radiat Biol 1984;11:295–389.

16. Suzuki M, Kase Y, Yamaguchi H, et al. Relative biological effectiveness for cell-killing effect on various human cell lines irradiated with heavy-ion medical accelerator in chiba (himac) carbon-ion beams. International journal of radiation oncology, biology, physics 2000;48:241–50.

17. Allalunis-Turner MJ, Barron GM, Day RS, 3rd, et al. Isolation of two cell lines from a human malignant glioma specimen differing in sensitivity to radiation and chemotherapeutic drugs. Radiat Res 1993;134:349–54.

18. Allalunis-Turner MJ, Zia PK, Barron GM, et al. Radiation-induced DNA damage and repair in cells of a radiosensitive human malignant glioma cell line. Radiat Res 1995;144:288–93.

19. Mitsudomi T, Steinberg SM, Nau MM, et al. P53 gene mutations in non-small-cell lung cancer cell lines and their correlation with the presence of ras mutations and clinical features. Oncogene 1992;7:171–80.

20. Friedrich T, Scholz U, Elsässer T, et al. Systematic analysis of rbe and related quantities using a database of cell survival experiments with ion beam irradiation. Journal of Radiation Research 2013;54:494–514.

21. Friedland W, Schmitt E, Kundrat P, et al. Comprehensive track-structure based evaluation of DNA damage by light ions from radiotherapy-relevant energies down to stopping. Sci Rep 2017;7:45161.

22. Luhr A, von Neubeck C, Helmbrecht S, et al. Modeling in vivo relative biological effectiveness in particle therapy for clinically relevant endpoints. Acta Oncol 2017;56:1392–1398.

23. Curtis SB. Lethal and potentially lethal lesions induced by radiation--a unified repair model. Radiat Res 1986;106:252–70.

24. Britten RA, Murray D. Constancy of the relative biological effectiveness of 42 mev (p-->be+) neutrons among cell lines with different DNA repair proficiencies. Radiat Res 1997;148:308–16.

25. Eguchi-Kasai K, Murakami M, Itsukaichi H, et al. Repair of DNA double-strand breaks and cell killing by charged particles. Adv Space Res 1998;22:543–9.

26. Takahashi A, Kubo M, Ma H, et al. Nonhomologous end-joining repair plays a more important role than homologous recombination repair in defining radiosensitivity after exposure to high-let radiation. Radiat Res 2014;182:338–44.

27. Karger CP, Peschke P. Rbe and related modeling in carbon-ion therapy. Phys Med Biol 2017;63:01TR02.

28. Hamada N, Hara T, Omura-Minamisawa M, et al. Energetic heavy ions overcome tumor radioresistance caused by overexpression of bcl-2. Radiotherapy and oncology : journal of the European Society for Therapeutic Radiology and Oncology 2008;89:231–6.

29. Jin XD, Gong L, Guo CL, et al. Survivin expressions in human hepatoma hepg2 cells exposed to ionizing radiation of different let. Radiat Environ Biophys 2008;47:399–404.

30. Krämer M, Scholz M. Treatment planning for heavy-ion radiotherapy: Calculation and optimization of biologically effective dose. Phys Med Biol 2000;45:3319–3330.

31. Scholz M, Kellerer AM, Kraft-Weyrather W, et al. Computation of cell survival in heavy ion beams for therapy. The model and its approximation. Radiat Environ Biophys 1997;36:59–66.

32. Damstrup L, Zimmerman A, Sirrenberg C, et al. M3814, a DNA-dependent protein kinase inhibitor (DNA-pki), potentiates the effect of ionizing radiation (ir) in xenotransplanted tumors in nude mice. International Journal of Radiation Oncology • Biology • Physics 2016;94:940–941.

33. Durant ST, Zheng L, Wang Y, et al. The brain-penetrant clinical atm inhibitor azd1390 radiosensitizes and improves survival of preclinical brain tumor models. Sci Adv 2018;4:eaat1719.

34. Leszczynska KB, Dobrynin G, Leslie RE, et al. Preclinical testing of an atr inhibitor demonstrates improved response to standard therapies for esophageal cancer. Radiotherapy and oncology : journal of the European Society for Therapeutic Radiology and Oncology 2016;121:232–238.

35. Grun R, Friedrich T, Traneus E, et al. Is the dose-averaged let a reliable predictor for the relative biological effectiveness? Medical physics 2019;46:1064–1074.

36. Kanai T, Furusawa Y, Fukutsu K, et al. Irradiation of mixed beam and design of spread-out bragg peak for heavy-ion radiotherapy. Radiat Res 1997;147:78–85.

