## Supplemental Material for "An empirical model to predict survival curve and relative biological effectiveness after helium and carbon ion irradiation based solely on the cell survival after photon irradiation"

### SUPPLEMENTAL INFORMATION

#### Supplemental Method 1: Cell Lines

We used a number of cell lines with differing sensitivities to photon radiation, genotypes, and originating from differing tumor sites (Supplemental Table 1).

Supplemental Table 1: Cell lines used in this work, their histology, and their intrinsic photon radiosensitivity ($D_{10\%,photon}$ - the dose for 10% survival).

| **Cell Line** | **Cancer Histology** | **$D_{10\%\ photons}$ (Gy)** |
|---|---|---|
| M059K | Glioblastoma | 5.41 ± 0.16 |
| M059J | Glioblastoma | 1.37 ± 0.02 |
| H1299 | Non-small cell lung cancer | 8.12 ± 0.08 |
| H460 | Large cell lung cancer | 4.81 ± 0.04 |
| BxPC-3 | Pancreatic Adenocarcenoma | 5.80 ± 0.12 |
| AsPC-1 | Pancreatic Adenocarcenoma | 5.38 ± 0.08 |
| PANC-1 | Pancreatic Adenocarcenoma | 6.85 ± 0.17 |
| Panc 10.05 | Pancreatic Adenocarcenoma | 8.56 ± 0.19 |

**Supplemental Method 2: Cell Culture Information**

Cells were cultured in humidified incubators at 37°C and 5% $CO_2$ at HIMAC and MDACC. The H460, H1299 and BxPC-3 cell lines were cultured in RPMI 1640 medium with 10% Fetal Bovine Serum (FBS) and 1% Penicillin/Streptomyicin (PS) added. The M059K and M059J cell lines were cultured in a 1:1 mixture of DMEM and F-12 Ham media with 10% FBS, 1% PS and 15 mM of HEPES buffer added. The AsPC-1, PANC-1 and Panc 10.05 cell lines were culture in high glucose DMEM with 10% FBS and 1% PS added. The technical details of the cell culture media and additives are given in Supplemental Table 2.

Prior to irradiation, the cells were cultured in T-25 and T-75 flasks before being seeded into either 6-well plates or T-12.5 flasks for the clonogenic assays. To detach the cells from the flasks, the cells were first washed with phosphate buffered saline (PBS), then treated with trypsin for 5 minutes. For the exposures at HIMAC, to ensure complete media coverage of the cells in the horizontal beamline, 2 hours prior to irradiation the 6-well plates were completely filled with ~15 ml of media and sealed with aluminum sealers, and the sealers were perforated immediately after irradiation to restore airflow. Details of these reagents and equipment can be found in Supplemental Table 3.

Supplemental Table 2: Cell culture media and additives and colony formation times.

| **Cell Line** | **Culture Medium (details)** | **Manufacturer (Cat. #)** | **Media Additives** | **Manufacturer (Cat. #)** | **Clonogenic Incubation Time** |
|---|---|---|---|---|---|
| M059K | 1:1 DMEM/F-12 Ham (with L-glutamine, $NaHCO_3$, without HEPES) | Sigma (D8062) | -10% FBS<br>-1% PS<br>-15 mM HEPES | -Sigma (F0926)<br>-Hyclone (SV30010)<br>-Sigma (H0887) | 9-10 days |
| M059J | | | | | 9-10 days |
| H1299 | RPMI 1640 (with L-glutamine, $NaHCO_3$) | Sigma (R8758) | -10% FBS<br>-1% PS | -Sigma (F0926)<br>-Hyclone (SV30010) | 8 days |
| H460 | | | | | 10 days |
| BxPC-3 | | | | | 14 days |
| AsPC-1 | DMEM – high glucose (with 4500 mg/L glucose, L-glutamine, sodium pyruvate, $NaHCO_3$) | Sigma (D6429) | -10% FBS<br>-1% PS | -Sigma (F0926)<br>-Hyclone (SV30010) | 14 days |
| PANC-1 | | | | | |
| Panc 10.05 | | | | | |

Supplemental Table 3: Reagents and equipment used to culture and process the cells.

| **Reagent/Equipment** | **Manufacturer (Cat. #)** | **Details** |
|---|---|---|
| T-25 flask | Thermo Scientific (156367) | Surface treated with Nunclon Delta, 25 $cm^2$ |
| T-75 flask | Thermo Scientific (156499) | Surface treated with Nunclon Delta, 75 $cm^2$ |
| T-12.5 flask | Cell Treat (229321) | 12.5 $cm^2$ |
| 6-well plates | Corning Costar (3506) | Treated for superior cell attachment |
| PBS | Hyclone (SH30256.01) | 0.0067 M ($PO_4$), with Ca and Mg |
| Trypsin | Corning (25-053-CL) | 0.25%, 2.21 mM EDTA, 1X $NaHCO_3$ |
| Aluminum Sealer | Bio Rad (MSF1001) | Free of DNase, RNase and DNA |
| Crystal Violet | Sigma (C0775-25G) | Dissolved in 100% ethanol |

**Supplemental Method 3: Irradiations**

The He- and C-ion irradiations were performed at HIMAC (Chiba, Japan) with pristine beams of nominal energy 150 MeV/nucleon (He-ions) and 290 MeV/nucleon (C-ions), using water-equivalent energy degraders to achieve dose-weighted LET values at the sample position of 2.20, 6.97 and 14.0 keV/µm for the He-ions, and 13.5, 27.9 and 60.5 keV/µm for the C-ions. All LET values used in this work were determined as summarized in Kanai et al., 1997 (36) and correspond to depths in water that are proximal to the Bragg peak for each beam. For a reference radiation condition, the cells were irradiated at The University of Texas MD Anderson Cancer Center with 6 MV photon from clinical linear accelerators at a water equivalent depth of 10 cm.

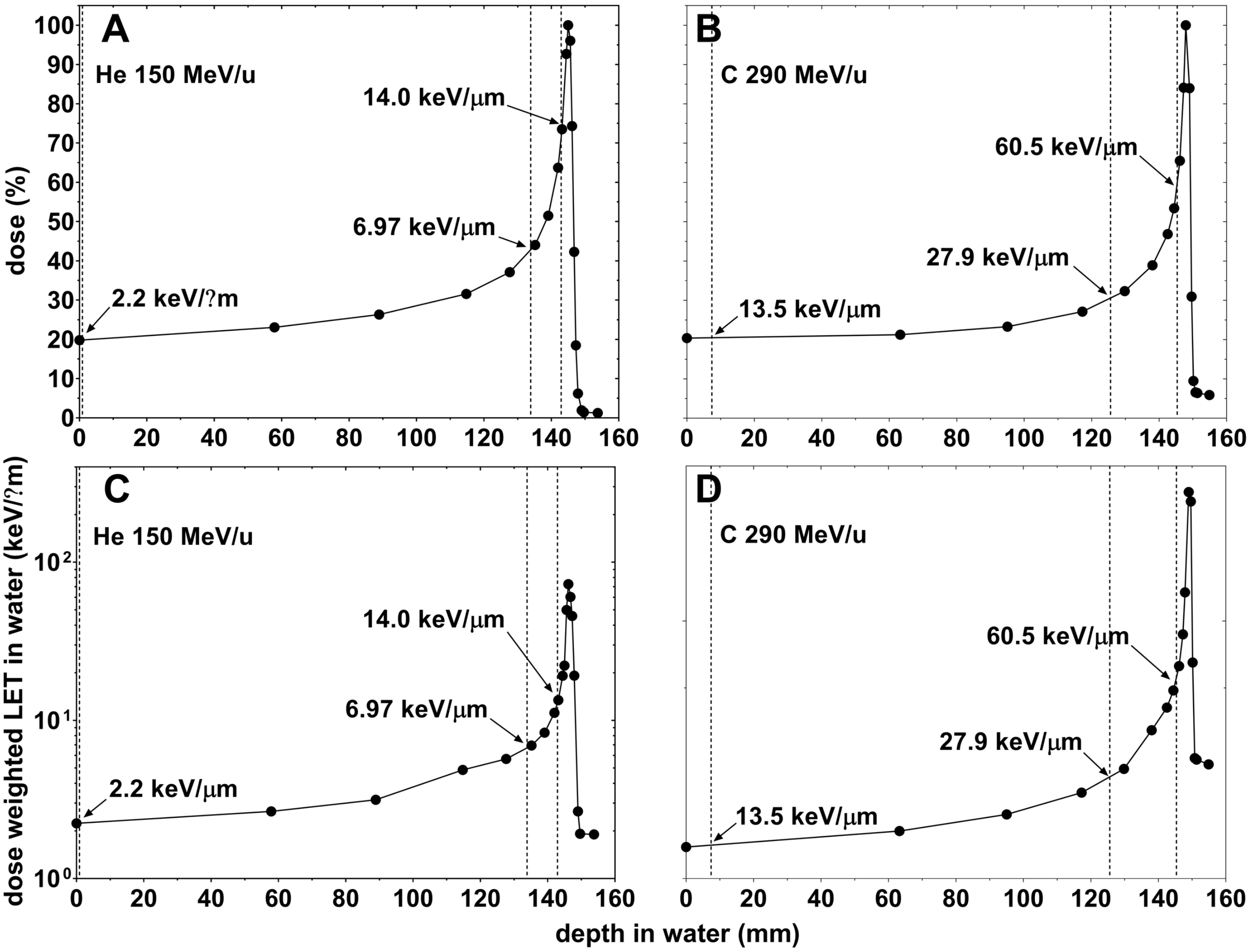


Supplemental Figure 1. Irradiation conditions for the He- and C-ion irradiations at HIMAC. Percentage depth dose for He- (A) and C-ions (B). Dose weighted LET in water for He- (C) and C-ions (D). Depths at which our cell lines were exposed are shown but a dashed line.

**Supplemental Method 4: Clonogenic Assays**

Cells were seeded into 6-well plates 24 hours before irradiation and exposed to doses of 0-8 Gy. After irradiation, the cells were incubated for 8-14 days (see Supplemental Table 2) before being fixed and stained with 0.5% crystal violet in ethanol solution. Colonies containing more than 50 cells were counted as viable. All experiments contained at least two replicates, and the experiments used to assess the LET trends in the data were independently repeated at least three times.

**Colony Counting**

The dishes containing the cells were scanned using an Epson Expression 10000 XL film scanner, and the images were evaluated using in-house ImageJ macros individually calibrated to score the colonies of each cell line. Briefly, these macros work by blurring the cells within a colony together using a Gaussian kernel whose radius corresponds to the radius of the smallest colony forming unit (CFU). Then contours are drawn around the blurred colonies, merged colonies are separated via watershedding, and colonies whose areas exceed the individually calibrated pixel area of the smallest CFU for that cell line are scored as colonies.

**Plating Efficiency Determination**

For each survival curve, the un-normalized survival fraction, i.e. the number of colonies counted divided by the number of wells seeded, was plotted as a function of dose. These data were fit to a function of the form:

$$SQ(D) = PE \cdot e^{-\alpha D - \beta D^2}$$

SQ is the survival quotient, and represents the un-normalized survival fraction, while the plating efficiency, PE, and α and β are free parameters in our fit. We fit this function using a variance-weighted least squares minimization to determine the value of PE and its uncertainty. This method results in less uncertainty in estimating the PE than simply using the survival quotient of the control dishes as it uses a larger data set to estimate the PE.

**Survival Fraction and Survival Curves**

For each experiment, i, we calculated the survival fraction $SF_{D,i}$ for each dose, D, after normalizing the survival quotient, $SQ_{D,i}$, by the plating efficiency $PE_i$:

$$SF_{D,i} = \frac{SQ_{D,i}}{PE_i}$$

Then, for each LET, a survival curve was produced by fitting all of the $SF_{D,i}$ simultaneously to the linear-quadratic (LQ) model:

$$SF_{D,i} = e^{-\alpha D - \beta D^2}$$

Where D is the dose in Gy, and α and β are free parameters determined by variance-weighted least-squares minimization. All the fitting was performed in Graphpad Prism 7.

**Supplemental Method 5: Drug treatment**

Ceralasertib (AZD6738, ATRi, Selleckchem) was dissolved in DMSO at 10 mM and was used at a final concentration of 0.1 μM. 8-(4-Dibenzothienyl)-2-(4-morpholinyl)-4H-1-benzopyran-4-one (NU7441, DNA-PKcsi, Selleckchem) was dissolved in DMSO at 5 mM and was used at 0.1 or 1 μM final concentrations. B02 (Cat# S8434, Rad51i, Selleckchem) was dissolved in DMSO at 10 mM and used at a 5 μM concentration. The final concentration of DMSO was 0.1% in all groups. Cells were seeded 24 hours prior to irradiation in 6 well plates, 8 hours prior to irradiation media was removed and replaced with media containing the inhibitor or DMSO vehicle. After 24 hours incubation with the inhibitor or vehicle, media was removed and replaced with fresh media. Cells were not washed to minimize disturbance to attached cells.

**Supplemental Method 6: Analyzing the PIDE Database**

Although the PIDE database has data for a hundreds of cell lines over a wide range of LET values (2-500 keV/μm for He- and C-ions), generally there aren't sufficient data at any particular LET to calculate the $R_{ion}$ versus $R_{photon}$ trends as we are able to do with our data. To work around this, we binned the data for similar LET values and fit those datasets as if they were the mean LET for that bin. And while this undoubtedly increases the uncertainty in these analyses, it nevertheless allows us to determine how the data trends for subsequent analyses. The bins themselves were chosen so as to cover a large range of LET values with the bins modestly spaced and containing a reasonable number of individual survival points. Details of the bins chosen are given in Supplemental Table 5.

As a final note, the PIDE database provides two sets of analyses that can be used: the α and β values of the linear quadratic model reported in the original publication or those calculated by the authors of the PIDE using the raw data from the publication. For consistency, we used the values calculated by the PIDE authors except where they were not provided or could not be used to determine all of the radiosensitivity parameters. A total of 49 LET values across 42 cell lines were used for He-ions and 113 LET values across 91 cell lines were used for C-ions.

Supplemental Table 4: Details of bins used to analyze the He-ion data in the PIDE database.

| Bin Number | Lowest LET in bin (keV/μm) | Highest LET in bin (keV/μm) | Mean LET in bin (keV/μm) | Standard Deviation of LETs in bin (keV/μm) | Number of data points in bin |
|---|---|---|---|---|---|
| 1 | 1.80 | 4.60 | 2.67 | 1.11 | 5 |
| 2 | 6.00 | 9.12 | 7.78 | 1.30 | 4 |
| 3 | 16.20 | 20.48 | 18.62 | 1.61 | 14 |
| 4 | 23.05 | 28.00 | 26.16 | 2.39 | 7 |
| 5 | 40.00 | 50.00 | 44.15 | 4.78 | 13 |
| 6 | 61.00 | 70.00 | 67.80 | 3.90 | 5 |
| 7 | 88.00 | 90.00 | 89.60 | 0.89 | 5 |
| 8 | 101.70 | 110.00 | 108.25 | 2.83 | 11 |
| 9 | 120.00 | 125.00 | 122.64 | 1.68 | 22 |

Supplemental Table 5: Details of bins used to analyze the C-ion data in the PIDE database.

| Bin Number | Lowest LET in bin (keV/μm) | Highest LET in bin (keV/μm) | Mean LET in bin (keV/μm) | Standard Deviation of LETs in bin (keV/μm) | Number of data points in bin |
|---|---|---|---|---|---|
| 1 | 12.16 | 13.80 | 13.14 | 0.39 | 41 |
| 2 | 19.00 | 23.00 | 20.60 | 1.13 | 21 |
| 3 | 29.50 | 33.60 | 30.83 | 1.27 | 22 |
| 4 | 38.00 | 40.60 | 39.65 | 0.69 | 29 |
| 5 | 46.00 | 50.30 | 49.58 | 1.20 | 25 |
| 6 | 55.00 | 61.50 | 59.26 | 2.02 | 8 |
| 7 | 68.00 | 70.00 | 69.90 | 0.45 | 20 |
| 8 | 76.00 | 77.10 | 76.93 | 0.36 | 20 |
| 9 | 78.50 | 81.00 | 80.25 | 0.63 | 19 |
| 10 | 88.00 | 94.00 | 92.10 | 2.60 | 10 |
| 11 | 99.00 | 103.00 | 100.77 | 1.42 | 13 |
| 12 | 105.00 | 110.00 | 107.17 | 2.04 | 12 |
| 13 | 150.00 | 153.50 | 151.75 | 1.63 | 12 |
| 14 | 220.00 | 225.00 | 221.88 | 1.55 | 8 |

**Supplemental Results 1: Summary of measured cell survival curvesSupplemental** Table 6 shows the survival parameters measured for each cell line at each radiation quality according to the linear quadratic model. The α and β values were determined in GraphPad7 via variance weighted least squares minimization, along with the normalized covariance of α and β ($NormCov(\alpha,\beta) = cov(\alpha,\beta)/\sigma_\alpha\sigma_\beta$). Then, the radiosensitivity metrics $D_{5\%}$, $D_{10\%}$, $D_{20\%}$, $D_{37\%}$, $D_{50\%}$ and $SF_{2Gy}$ were calculated, along with their uncertainties, from the values of α, β and NormCov(α,β).

Supplemental Table 6: Summary of survival parameters measured for our cell line panel at all radiation qualities: α and β from the linear quadratic model, their normalized covariance, NormCov(α, β), the dose for 5%, 10%, 20%, 37% and 50% survival ($D_{5\%}$, $D_{10\%}$, $D_{20\%}$, $D_{37\%}$, and $D_{50\%}$), the mean inactivation dose, $\bar{D}$, the surviving fraction for a dose of 2 Gy, $SF_{2Gy}$, and for the ion conditions, the RBE using $D_{10\%}$ as the biological endpoint, $RBE_{D10\%}$. For some survival curves the linear quadratic model gave a very small β value. In those cases, β was arbitrarily set to $10^{-5}$ $Gy^{-2}$ to allow the calculations of the parameters of the survival curve.

| Radiation type | Cell line | α ($Gy^{-1}$) | β ($Gy^{-2}$) | Norm cov(α,β) | $D_{5\%}$ (Gy) | $D_{10\%}$ (Gy) | $D_{20\%}$ (Gy) | $D_{37\%}$ (Gy) | $D_{50\%}$ (Gy) | $SF_{2Gy}$ (Gy) | $\bar{D}$ (Gy) | $RBE_{D(10\%)}$ |
|---|---|---|---|---|---|---|---|---|---|---|---|---|
| 6 MV Photons | H460 | 0.046±0.033 | 0.0899±0.0066 | -0.9729 | 5.523±0.047 | 4.812±0.040 | 3.983±0.050 | 3.080±0.072 | 2.533±0.087 | 0.637±0.026 | 2.716±0.185 | 1.000±0.012 |
| | H1299 | 0.089±0.018 | 0.0240±0.0024 | -0.9746 | 9.475±0.115 | 8.119±0.076 | 6.546±0.074 | 4.848±0.111 | 3.834±0.133 | 0.761±0.021 | 4.256±0.285 | 1.000±0.013 |
| | M059K | 0.314±0.055 | 0.0206±0.0117 | -0.9724 | 6.639±0.309 | 5.407±0.162 | 4.046±0.120 | 2.689±0.162 | 1.954±0.165 | 0.491±0.032 | 2.488±0.374 | 1.000±0.042 |
| | M059J | 1.360±0.107 | 0.2318±0.0717 | -0.9617 | 1.706±0.028 | 1.372±0.020 | 1.010±0.023 | 0.657±0.025 | 0.472±0.022 | 0.026±0.003 | 0.619±0.043 | 1.000±0.021 |
| | BxPC-3 | 0.308±0.029 | 0.0154±0.0042 | -0.9477 | 7.162±0.131 | 5.797±0.117 | 4.301±0.130 | 2.829±0.131 | 2.042±0.117 | 0.508±0.022 | 2.641±0.207 | 1.000±0.029 |
| | AsPC-1 | 0.134±0.037 | 0.0546±0.0071 | -0.9613 | 6.282±0.106 | 5.383±0.079 | 4.341±0.079 | 3.215±0.107 | 2.543±0.124 | 0.615±0.029 | 2.822±0.249 | 1.000±0.021 |
| | PANC-1 | 0.136±0.045 | 0.0292±0.0074 | -0.9705 | 8.064±0.260 | 6.851±0.167 | 5.451±0.135 | 3.953±0.192 | 3.070±0.227 | 0.678±0.041 | 3.484±0.495 | 1.000±0.034 |
| | Panc 10.05 | 0.113±0.024 | 0.0182±0.0034 | -0.9528 | 10.096±0.279 | 8.563±0.190 | 6.797±0.139 | 4.911±0.166 | 3.802±0.191 | 0.741±0.027 | 4.333±0.426 | 1.000±0.031 |
| | H460+DMSO | 0.013±0.022 | 0.0790±0.0043 | -0.9723 | 6.076±0.044 | 5.316±0.033 | 4.431±0.034 | 3.465±0.050 | 2.880±0.062 | 0.710±0.019 | 3.071±0.155 | 1.000±0.009 |
| | H460+NU7441 (DNA-PKcsi, 0.1 μM) | 0.237±0.069 | 0.1125±0.0160 | -0.9586 | 4.214±0.070 | 3.592±0.075 | 2.874±0.093 | 2.101±0.113 | 1.644±0.121 | 0.397±0.032 | 1.848±0.195 | 1.000±0.029 |
| | H460+AZD6738 (ATRi, 0.1 μM) | 0.269±0.053 | 0.0963±0.0128 | -0.9602 | 4.352±0.061 | 3.687±0.058 | 2.922±0.071 | 2.106±0.088 | 1.627±0.094 | 0.397±0.023 | 1.860±0.158 | 1.000±0.022 |
| | H1299+DMSO | 0.077±0.023 | 0.0204±0.0029 | -0.9754 | 10.383±0.196 | 8.908±0.126 | 7.198±0.101 | 5.348±0.151 | 4.242±0.188 | 0.790±0.027 | 4.694±0.435 | 1.000±0.020 |
| | H1299+B02 (Rad51i, 0.5 μM) | 0.127±0.029 | 0.0158±0.0037 | -0.9694 | 10.330±0.278 | 8.707±0.176 | 6.845±0.153 | 4.873±0.215 | 3.727±0.244 | 0.728±0.032 | 4.322±0.506 | 1.000±0.029 |
| | H1299+NU7441 (DNA-PKcsi, 0.1 μM) | 0.159±0.025 | 0.0218±0.0040 | -0.9549 | 8.621±0.202 | 7.250±0.134 | 5.681±0.108 | 4.023±0.135 | 3.064±0.149 | 0.666±0.024 | 3.576±0.311 | 1.000±0.026 |
| | H1299+NU7441 (DNA-PKcsi, 1 μM) | 1.068±0.122 | 0.0000±0.0123 | -0.9296 | 2.805±0.238 | 2.156±0.197 | 1.507±0.148 | 0.931±0.097 | 0.649±0.070 | 0.118±0.023 | 0.936±0.107 | 1.000±0.129 |
| | H1299+AZD6738 (ATRi, 0.1 μM) | 0.259±0.032 | 0.0040±0.0051 | -0.9547 | 10.006±0.646 | 7.914±0.325 | 5.706±0.184 | 3.632±0.199 | 2.572±0.186 | 0.586±0.027 | 3.506±0.508 | 1.000±0.058 |
| | PANC-1+DMSO | 0.182±0.038 | 0.0297±0.0058 | -0.9636 | 7.437±0.142 | 6.259±0.115 | 4.909±0.136 | 3.482±0.174 | 2.656±0.185 | 0.617±0.033 | 3.093±0.328 | 1.000±0.026 |
| | Panc1+NU7441 (DNA-PKcsi, 0.1 μM) | 0.154±0.041 | 0.0421±0.0064 | -0.9582 | 6.798±0.117 | 5.785±0.109 | 4.615±0.132 | 3.360±0.166 | 2.619±0.179 | 0.621±0.036 | 2.959±0.305 | 1.000±0.027 |
| | Panc1+AZD6738 (ATRi, 0.1 μM) | 0.143±0.038 | 0.0362±0.0058 | -0.9629 | 7.334±0.128 | 6.242±0.109 | 4.980±0.130 | 3.627±0.170 | 2.827±0.188 | 0.650±0.036 | 3.193±0.335 | 1.000±0.025 |
| | Panc1005+DMSO | 0.214±0.050 | 0.0129±0.0066 | -0.9645 | 9.047±0.361 | 7.424±0.241 | 5.612±0.264 | 3.778±0.319 | 2.771±0.313 | 0.619±0.046 | 3.460±0.601 | 1.000±0.046 |
| | Panc1005+NU7441 (DNA-PKcsi, 0.1 μM) | 0.282±0.034 | 0.0164±0.0042 | -0.9594 | 7.421±0.137 | 6.041±0.150 | 4.518±0.174 | 3.000±0.174 | 2.180±0.156 | 0.533±0.028 | 2.777±0.254 | 1.000±0.035 |
| | Panc1005+AZD6738 (ATRi, 0.1 μM) | 0.269±0.071 | 0.0228±0.0095 | -0.9561 | 6.993±0.248 | 5.754±0.267 | 4.367±0.313 | 2.955±0.329 | 2.175±0.305 | 0.533±0.057 | 2.696±0.483 | 1.000±0.066 |
| 2.2 keV/μm He-ions | H460 | 0.182±0.055 | 0.0821±0.0108 | -0.9653 | 5.033±0.072 | 4.302±0.075 | 3.455±0.096 | 2.543±0.123 | 2.001±0.134 | 0.500±0.034 | 2.234±0.223 | 1.119±0.022 |
| | H1299 | 0.230±0.052 | 0.0105±0.0079 | -0.9689 | 9.194±0.561 | 7.479±0.301 | 5.587±0.232 | 3.705±0.300 | 2.690±0.301 | 0.606±0.045 | 3.434±0.675 | 1.086±0.045 |
| | M059K | 0.112±0.018 | 0.0709±0.0036 | -0.9569 | 5.762±0.038 | 4.967±0.032 | 4.043±0.035 | 3.041±0.046 | 2.438±0.053 | 0.603±0.014 | 2.668±0.104 | 1.089±0.033 |
| | M059J | 1.034±0.163 | 0.5182±0.1155 | -0.9674 | 1.605±0.030 | 1.334±0.023 | 1.027±0.028 | 0.709±0.035 | 0.530±0.035 | 0.016±0.003 | 0.639±0.063 | 1.028±0.023 |
| | BxPC3 | 0.319±0.062 | 0.0090±0.0112 | -0.9681 | 7.715±0.510 | 6.153±0.254 | 4.481±0.209 | 2.884±0.250 | 2.055±0.234 | 0.510±0.042 | 2.750±0.552 | 0.942±0.043 |
| 7.0 keV/μm He-ions | H460 | 0.122±0.048 | 0.1050±0.0123 | -0.9708 | 4.792±0.073 | 4.138±0.052 | 3.377±0.050 | 2.551±0.073 | 2.053±0.089 | 0.515±0.025 | 2.239±0.196 | 1.163±0.017 |
| | H1299 | 0.197±0.041 | 0.0163±0.0059 | -0.9721 | 8.798±0.279 | 7.289±0.169 | 5.586±0.172 | 3.832±0.232 | 2.847±0.243 | 0.632±0.037 | 3.464±0.483 | 1.114±0.028 |
| | M059K | 0.364±0.040 | 0.0242±0.0080 | -0.9591 | 5.912±0.130 | 4.799±0.091 | 3.575±0.100 | 2.362±0.112 | 1.711±0.104 | 0.439±0.022 | 2.196±0.201 | 1.127±0.040 |
| | M059J | 1.425±0.301 | 0.2688±0.2121 | -0.9739 | 1.612±0.059 | 1.298±0.043 | 0.957±0.056 | 0.624±0.062 | 0.448±0.056 | 0.020±0.006 | 0.587±0.107 | 1.057±0.039 |
| | BxPC3 | 0.361±0.089 | 0.0130±0.0148 | -0.9667 | 6.689±0.328 | 5.349±0.253 | 3.908±0.305 | 2.524±0.317 | 1.803±0.280 | 0.461±0.056 | 2.398±0.549 | 1.084±0.056 |
| 14.0 keV/μm He-ions | H460 | 0.089±0.056 | 0.1328±0.0159 | -0.9734 | 4.426±0.071 | 3.842±0.050 | 3.162±0.047 | 2.421±0.069 | 1.973±0.086 | 0.492±0.026 | 2.130±0.205 | 1.253±0.019 |
| | H1299 | 0.153±0.030 | 0.0287±0.0048 | -0.9483 | 7.898±0.173 | 6.684±0.126 | 5.286±0.115 | 3.798±0.142 | 2.927±0.156 | 0.657±0.028 | 3.357±0.305 | 1.215±0.026 |
| | M059K | 0.553±0.059 | 0.0000±0.0160 | -0.9535 | 5.419±0.575 | 4.165±0.442 | 2.911±0.309 | 1.799±0.191 | 1.254±0.133 | 0.331±0.020 | 1.809±0.192 | 1.298±0.143 |
| | M059J | 1.326±0.110 | 0.4942±0.0922 | -0.9702 | 1.462±0.020 | 1.200±0.013 | 0.907±0.014 | 0.611±0.018 | 0.448±0.018 | 0.010±0.002 | 0.559±0.035 | 1.144±0.021 |
| | BxPC3 | 0.453±0.073 | 0.0102±0.0141 | -0.9365 | 5.847±0.297 | 4.608±0.218 | 3.309±0.215 | 2.098±0.198 | 1.482±0.166 | 0.388±0.037 | 2.035±0.344 | 1.258±0.065 |
| | H1299+DMSO | 0.196±0.016 | 0.0175±0.0022 | -0.9787 | 8.623±0.077 | 7.157±0.052 | 5.499±0.070 | 3.785±0.093 | 2.820±0.096 | 0.629±0.015 | 3.415±0.178 | 1.245±0.020 |
| | H1299+B02 (Rad51i, 5 μM) | 0.235±0.026 | 0.0276±0.0046 | -0.9612 | 6.998±0.115 | 5.821±0.078 | 4.486±0.078 | 3.102±0.099 | 2.318±0.103 | 0.560±0.019 | 2.791±0.197 | 1.496±0.036 |
| | H1299+NU7441 (DNA-PKcsi, 1 μM) | 1.591±0.267 | 0.0000±0.0409 | -0.8995 | 1.883±0.237 | 1.447±0.196 | 1.012±0.146 | 0.625±0.096 | 0.436±0.069 | 0.042±0.016 | 0.629±0.105 | 1.490±0.243 |
| 13.5 keV/μm C-ions | H460 | 0.307±0.096 | 0.0740±0.0291 | -0.9723 | 4.621±0.218 | 3.879±0.131 | 3.032±0.092 | 2.139±0.132 | 1.624±0.152 | 0.403±0.034 | 1.905±0.338 | 1.240±0.043 |
| | H1299 | 0.045±0.029 | 0.0485±0.0048 | -0.9412 | 7.412±0.124 | 6.445±0.101 | 5.318±0.095 | 4.089±0.117 | 3.346±0.136 | 0.753±0.031 | 3.601±0.298 | 1.260±0.023 |
| | M059K | 0.206±0.061 | 0.0886±0.0194 | -0.9691 | 4.768±0.166 | 4.067±0.109 | 3.256±0.065 | 2.384±0.075 | 1.867±0.095 | 0.465±0.023 | 2.097±0.251 | 1.329±0.054 |
| | M059J | 0.944±0.389 | 0.7848±0.3312 | -0.9652 | 1.443±0.065 | 1.214±0.045 | 0.952±0.046 | 0.675±0.062 | 0.514±0.067 | 0.007±0.004 | 0.600±0.122 | 1.130±0.045 |
| | BxPC3 | 0.458±0.064 | 0.0002±0.0167 | -0.9699 | 6.527±0.705 | 5.020±0.295 | 3.512±0.121 | 2.171±0.141 | 1.514±0.131 | 0.400±0.026 | 2.181±0.458 | 1.155±0.072 |

| | | | | | | | | | | | | |
|---|---|---|---|---|---|---|---|---|---|---|---|---|
| | AsPC-1 | 0.143±0.036 | 0.0709±0.0074 | -0.9716 | 5.570±0.064 | 4.780±0.050 | 3.862±0.059 | 2.870±0.084 | 2.277±0.098 | 0.566±0.025 | 2.519±0.190 | 1.126±0.020 |
| | PANC-1 | 0.227±0.066 | 0.0256±0.0123 | -0.978 | 7.250±0.346 | 6.029±0.191 | 4.645±0.153 | 3.211±0.231 | 2.399±0.255 | 0.573±0.048 | 2.889±0.549 | 1.136±0.045 |
| | Panc1005 | 0.151±0.041 | 0.0280±0.0058 | -0.9749 | 7.999±0.153 | 6.770±0.116 | 5.356±0.147 | 3.849±0.205 | 2.967±0.226 | 0.662±0.039 | 3.402±0.416 | 1.265±0.035 |
| | H460+DMSO | 0.000±0.041 | 0.1165±0.0120 | -0.9587 | 5.071±0.105 | 4.446±0.078 | 3.717±0.055 | 2.921±0.053 | 2.439±0.065 | 0.628±0.024 | 2.596±0.220 | 1.196±0.022 |
| | H460+NU7441 (DNA-PKcsi, 0.1 µM) | 0.030±0.014 | 0.1224±0.0040 | -0.9735 | 4.828±0.026 | 4.218±0.018 | 3.507±0.013 | 2.732±0.016 | 2.262±0.021 | 0.578±0.007 | 2.417±0.065 | 0.852±0.018 |
| | H460+AZD6738 (ATRi, 0.1 µM) | 0.103±0.059 | 0.1268±0.0190 | -0.9689 | 4.471±0.115 | 3.874±0.080 | 3.179±0.052 | 2.423±0.061 | 1.966±0.079 | 0.490±0.023 | 2.129±0.225 | 0.952±0.025 |
| | H1299+DMSO | 0.033±0.057 | 0.0478±0.0083 | -0.9655 | 7.574±0.168 | 6.599±0.148 | 5.464±0.176 | 4.225±0.243 | 3.475±0.289 | 0.773±0.064 | 3.727±0.604 | 1.350±0.036 |
| | H1299+NU7441 (DNA-PKcsi, 0.1 µM) | 0.126±0.016 | 0.0427±0.0030 | -0.9757 | 7.032±0.067 | 6.017±0.042 | 4.840±0.032 | 3.571±0.047 | 2.816±0.059 | 0.655±0.013 | 3.137±0.137 | 1.205±0.024 |
| | H1299+AZD6738 (ATRi, 0.1 µM) | 0.062±0.029 | 0.0425±0.0047 | -0.9767 | 7.699±0.111 | 6.668±0.075 | 5.469±0.068 | 4.164±0.104 | 3.377±0.132 | 0.746±0.029 | 3.659±0.313 | 1.187±0.051 |
| | Panc1+DMSO | 0.117±0.024 | 0.0426±0.0045 | -0.98 | 7.127±0.096 | 6.108±0.059 | 4.927±0.045 | 3.651±0.071 | 2.889±0.090 | 0.668±0.020 | 3.205±0.212 | 1.025±0.021 |
| | Panc1+NU7441 (DNA-PKcsi, 0.1 µM) | 0.124±0.015 | 0.0478±0.0031 | -0.9807 | 6.732±0.056 | 5.770±0.035 | 4.654±0.025 | 3.449±0.040 | 2.730±0.051 | 0.645±0.012 | 3.028±0.122 | 1.003±0.020 |
| | Panc1+AZD6738 (ATRi, 0.1 µM) | 0.174±0.026 | 0.0324±0.0053 | -0.9761 | 7.300±0.169 | 6.164±0.103 | 4.858±0.056 | 3.471±0.074 | 2.663±0.093 | 0.620±0.019 | 3.074±0.237 | 1.013±0.025 |
| | Panc1005+DMSO | 0.345±0.007 | 0.0000±0.0000 | -0.7559 | 8.674±0.177 | 6.667±0.136 | 4.660±0.095 | 2.879±0.059 | 2.007±0.041 | 0.501±0.007 | 2.896±0.060 | 1.113±0.043 |
| | Panc1005+NU7441 (DNA-PKcsi, 0.1 µM) | 0.424±0.064 | 0.0000±0.0074 | -0.9387 | 7.067±1.066 | 5.432±0.819 | 3.797±0.573 | 2.345±0.354 | 1.635±0.247 | 0.428±0.043 | 2.359±0.356 | 1.112±0.170 |
| | Panc1005+AZD6738 (ATRi, 0.1 µM) | 0.138±0.091 | 0.0809±0.0288 | -0.9662 | 5.290±0.364 | 4.548±0.255 | 3.687±0.149 | 2.754±0.118 | 2.195±0.152 | 0.549±0.042 | 2.416±0.497 | 1.265±0.092 |
| | H460 | 0.502±0.048 | 0.0892±0.0174 | -0.9773 | 3.627±0.060 | 2.993±0.033 | 2.280±0.031 | 1.551±0.044 | 1.146±0.047 | 0.256±0.008 | 1.410±0.097 | 1.608±0.022 |
| | H1299 | 0.098±0.070 | 0.0748±0.0139 | -0.972 | 5.707±0.120 | 4.932±0.097 | 4.029±0.117 | 3.048±0.167 | 2.458±0.199 | 0.609±0.053 | 2.676±0.401 | 1.646±0.036 |
| | M059K | 0.649±0.101 | 0.0000±0.0000 | -0.9684 | 4.616±0.720 | 3.548±0.553 | 2.480±0.387 | 1.532±0.239 | 1.068±0.167 | 0.273±0.055 | 1.541±0.261 | 1.524±0.242 |
| 27.9 keV/µm C-ions | M059J | 1.823±0.198 | 0.2446±0.1673 | -0.9648 | 1.386±0.037 | 1.101±0.024 | 0.798±0.028 | 0.510±0.029 | 0.363±0.025 | 0.010±0.003 | 0.490±0.052 | 1.247±0.033 |
| | BxPC3 | 0.543±0.126 | 0.0102±0.0373 | -0.9803 | 5.039±0.538 | 3.947±0.217 | 2.815±0.146 | 1.772±0.192 | 1.247±0.178 | 0.324±0.036 | 1.734±0.491 | 1.469±0.086 |
| | AsPC-1 | 0.361±0.107 | 0.0802±0.0271 | -0.974 | 4.263±0.109 | 3.561±0.098 | 2.763±0.132 | 1.928±0.166 | 1.452±0.170 | 0.352±0.039 | 1.726±0.292 | 1.512±0.047 |
| | PANC-1 | 0.196±0.033 | 0.0553±0.0071 | -0.9693 | 5.801±0.082 | 4.922±0.057 | 3.909±0.060 | 2.825±0.083 | 2.189±0.095 | 0.542±0.022 | 2.493±0.185 | 1.392±0.038 |
| | Panc1005 | 0.202±0.065 | 0.0342±0.0108 | -0.9718 | 6.859±0.191 | 5.766±0.148 | 4.515±0.187 | 3.194±0.248 | 2.431±0.264 | 0.582±0.052 | 2.840±0.475 | 1.485±0.050 |
| | H460 | 0.676±0.173 | 0.0845±0.0797 | -0.9743 | 3.173±0.242 | 2.576±0.124 | 1.920±0.081 | 1.269±0.111 | 0.920±0.114 | 0.184±0.015 | 1.179±0.269 | 1.868±0.091 |
| | H1299 | 0.124±0.048 | 0.1020±0.0124 | -0.9714 | 4.846±0.076 | 4.182±0.054 | 3.411±0.052 | 2.573±0.075 | 2.069±0.092 | 0.519±0.026 | 2.258±0.204 | 1.941±0.031 |
| | M059K | 0.765±0.036 | 0.0000±0.0283 | -0.9624 | 3.914±0.390 | 3.008±0.200 | 2.103±0.072 | 1.299±0.017 | 0.906±0.016 | 0.216±0.010 | 1.307±0.062 | 1.797±0.131 |
| | M059J | 1.365±0.442 | 0.8363±0.4291 | -0.9627 | 1.245±0.058 | 1.033±0.040 | 0.793±0.043 | 0.546±0.054 | 0.407±0.055 | 0.002±0.002 | 0.493±0.095 | 1.328±0.055 |
| | BxPC3 | 0.740±0.157 | 0.0000±0.0812 | -0.9682 | 4.051±0.991 | 3.114±0.454 | 2.176±0.136 | 1.344±0.106 | 0.937±0.109 | 0.228±0.019 | 1.352±0.288 | 1.862±0.274 |
| | AsPC-1 | 0.703±0.035 | 0.0000±0.0030 | -0.8257 | 4.261±0.157 | 3.275±0.130 | 2.289±0.098 | 1.414±0.065 | 0.986±0.046 | 0.245±0.015 | 1.422±0.072 | 1.644±0.070 |
| | PANC-1 | 0.197±0.072 | 0.0924±0.0161 | -0.9699 | 4.728±0.082 | 4.039±0.078 | 3.242±0.101 | 2.384±0.135 | 1.874±0.150 | 0.466±0.039 | 2.095±0.262 | 1.696±0.053 |
| | Panc1005 | 0.396±0.048 | 0.0415±0.0108 | -0.9754 | 4.971±0.076 | 4.073±0.062 | 3.072±0.084 | 2.063±0.099 | 1.510±0.095 | 0.383±0.021 | 1.893±0.172 | 2.103±0.057 |
| | H460+DMSO | 0.248±0.042 | 0.1975±0.0163 | -0.9653 | 3.317±0.037 | 2.844±0.026 | 2.295±0.023 | 1.702±0.031 | 1.348±0.037 | 0.276±0.008 | 1.494±0.081 | 1.869±0.021 |
| 60.5 keV/µm C-ions | H460+NU7441 (DNA-PKcsi, 0. 1 µM) | 0.335±0.052 | 0.1937±0.0229 | -0.9745 | 3.163±0.050 | 2.691±0.031 | 2.146±0.022 | 1.561±0.032 | 1.216±0.039 | 0.236±0.006 | 1.375±0.090 | 1.335±0.032 |
| | H460+AZD6738 (ATRi, 0.1 µM) | 0.596±0.214 | 0.1535±0.0960 | -0.9759 | 2.883±0.161 | 2.390±0.092 | 1.833±0.090 | 1.259±0.127 | 0.936±0.135 | 0.164±0.016 | 1.137±0.273 | 1.543±0.064 |
| | H1299+DMSO | 0.134±0.033 | 0.1018±0.0078 | -0.9566 | 4.807±0.049 | 4.144±0.041 | 3.373±0.044 | 2.536±0.059 | 2.034±0.068 | 0.509±0.019 | 2.226±0.132 | 2.150±0.037 |
| | H1299+NU7441 (DNA-PKcsi, 0.1 µM) | 0.241±0.047 | 0.0933±0.0136 | -0.968 | 4.521±0.082 | 3.842±0.054 | 3.059±0.047 | 2.220±0.066 | 1.725±0.077 | 0.425±0.019 | 1.956±0.163 | 1.887±0.044 |
| | H1299+AZD6738 (ATRi, 0.1 µM] | 0.123±0.067 | 0.0986±0.0189 | -0.9686 | 4.923±0.148 | 4.248±0.101 | 3.464±0.072 | 2.612±0.093 | 2.100±0.118 | 0.527±0.033 | 2.293±0.301 | 1.863±0.088 |
| | Panc1+DMSO | 0.175±0.050 | 0.1064±0.0113 | -0.9658 | 4.549±0.053 | 3.903±0.052 | 3.154±0.067 | 2.345±0.087 | 1.861±0.097 | 0.461±0.026 | 2.058±0.170 | 1.603±0.036 |
| | Panc1+NU7441 (DNA-PKcsi, 0.1 µM) | 0.159±0.032 | 0.1239±0.0089 | -0.9746 | 4.317±0.037 | 3.717±0.026 | 3.019±0.027 | 2.263±0.040 | 1.809±0.048 | 0.443±0.013 | 1.986±0.103 | 1.556±0.031 |
| | Panc1+AZD6738 (ATRi, 0.1 µM) | 0.329±0.050 | 0.0798±0.0143 | -0.9745 | 4.404±0.078 | 3.693±0.048 | 2.881±0.049 | 2.027±0.070 | 1.536±0.077 | 0.376±0.017 | 1.807±0.154 | 1.690±0.037 |
| | Panc1005+DMSO | 0.567±0.099 | 0.0252±0.0232 | -0.9612 | 4.419±0.159 | 3.515±0.145 | 2.551±0.167 | 1.636±0.161 | 1.163±0.137 | 0.291±0.033 | 1.566±0.261 | 2.112±0.111 |
| | Panc1005+NU7441 (DNA-PKcsi, 0.1 µM) | 0.302±0.238 | 0.1638±0.0601 | -0.9916 | 3.454±0.101 | 2.940±0.155 | 2.346±0.219 | 1.709±0.271 | 1.333±0.287 | 0.284±0.068 | 1.505±0.435 | 2.055±0.120 |
| | Panc1005+AZD6738 (ATRi, 0.1 µM) | 0.798±0.129 | 0.0889±0.0385 | -0.9643 | 2.850±0.081 | 2.298±0.095 | 1.697±0.106 | 1.109±0.099 | 0.798±0.085 | 0.142±0.017 | 1.041±0.137 | 2.504±0.155 |

**Supplemental Results 2: The linear correlations between radiosensitivity metrics are stronger for parameters corresponding to lower survival levels**

We calculated the radiosensitivity metrics $D_{5\%}$, $D_{10\%}$, $D_{20\%}$, $D_{37\%}$, $D_{50\%}$ and $SF_{2Gy}$ and saw that independent of the parameter used to characterize radiosensitivity, radiosensitivity to photons and ions are linearly related, but to differing degrees. Supplemental Figure 2 shows the correlations for the parameters that are not shown in Fig. 1:

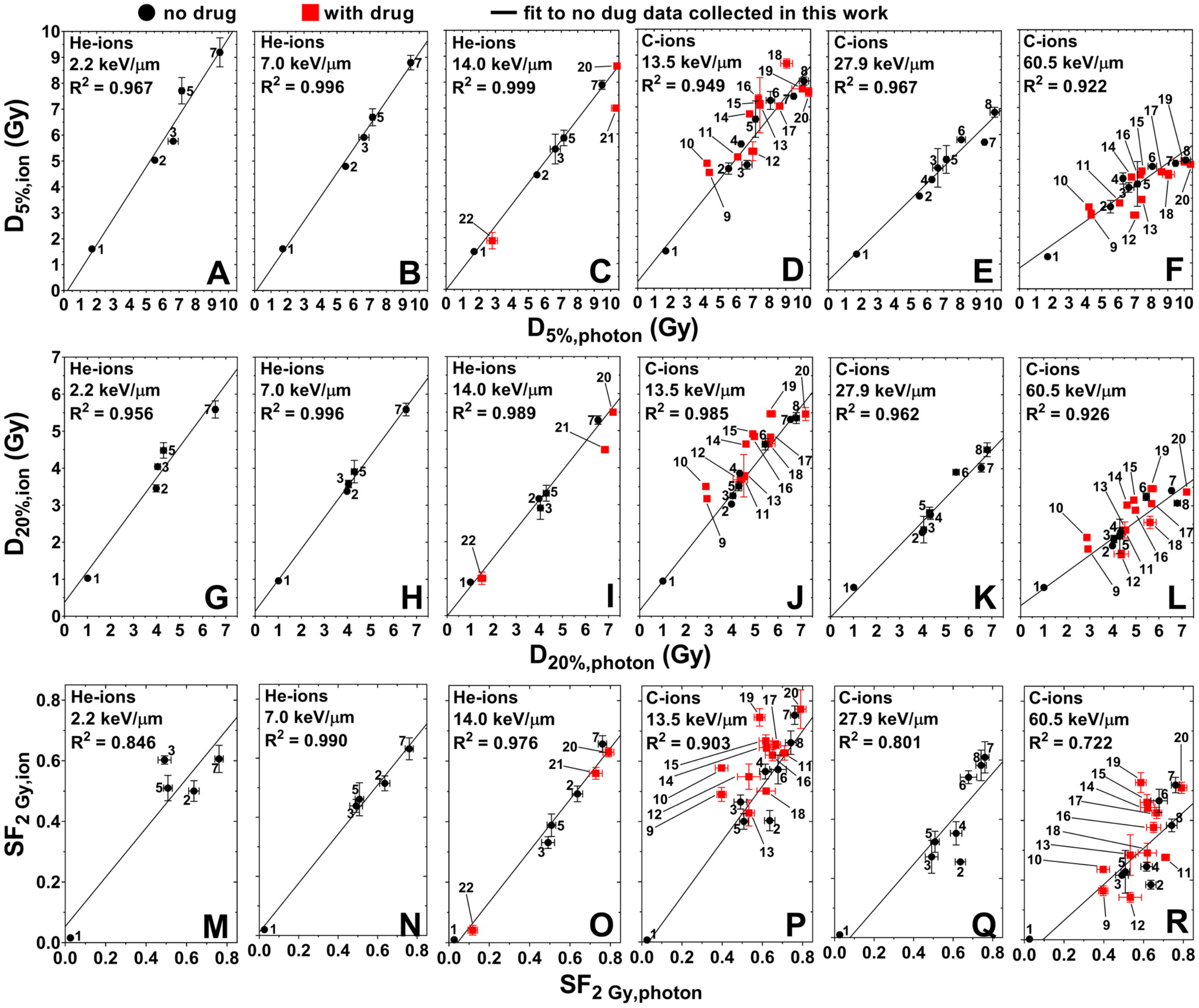


Supplemental Figure 2: (A-F) $D_{5\%,ion}$ vs. $D_{5\%,photon}$, (G-L) $D_{20\%,ion}$ vs. $D_{20\%,photon}$, and (M-R) $SF_{2Gy,ion}$ vs. $SF_{2Gy,photon}$ for He- and C-ions in order of increasing LET from 2.2 to 60.5 keV/µm. Black circles and red squares represent data of cell lines treated with radiation alone and drug in combination of radiation. Lines represent linear fits to the data of cell lines exposed to radiation alone.

Supplemental Table 7: $R^2$ values determined for the correlations between ion and photon radiosensitivity for the metrics $D_{5\%}$ , $D_{10\%}$ , $D_{20\%}$,, $D_{37\%}$,, $D_{50\%}$, and $SF_{2Gy}$. Abbreviations – LET: linear energy transfer; $D_{5\%}$ $D_{10\%}$, … $D_{50\%}$: dose for 5%, 10%, …, 50% survival; $SF_{2Gy}$: surviving fraction for a dose of 2 Gy.

| Ion | LET (keV/µm) | $R^2$ value | | | | | |
|---|---|---|---|---|---|---|---|
| | | $D_{5\%}$ | $D_{10\%}$ | $D_{20\%}$ | $D_{37\%}$ | $D_{50\%}$ | $SF_{2Gy}$ |
| He-ions | 2.2 | 0.9665 | 0.9732 | 0.9559 | 0.8722 | 0.7621 | 0.8464 |
| | 7.0 | 0.9956 | 0.9968 | 0.9963 | 0.9912 | 0.9837 | 0.9903 |
| | 14.0 | 0.9986 | 0.9954 | 0.9886 | 0.9822 | 0.981 | 0.9762 |
| C-ions | 13.5 | 0.9489 | 0.9737 | 0.9847 | 0.9638 | 0.9248 | 0.9031 |
| | 27.9 | 0.9667 | 0.9714 | 0.9621 | 0.9448 | 0.9118 | 0.8008 |
| | 60.5 | 0.9224 | 0.9462 | 0.9261 | 0.8516 | 0.7824 | 0.7215 |

**Supplemental Results 3: Summary PIDE database fits**

For the all the radiosensitivity parameters, we noted that the slope and intercept of the correlations varied regularly with LET. Supplemental Figure 3 shows the trends for the radiosensitivity parameters that were not shown in Fig. 2:

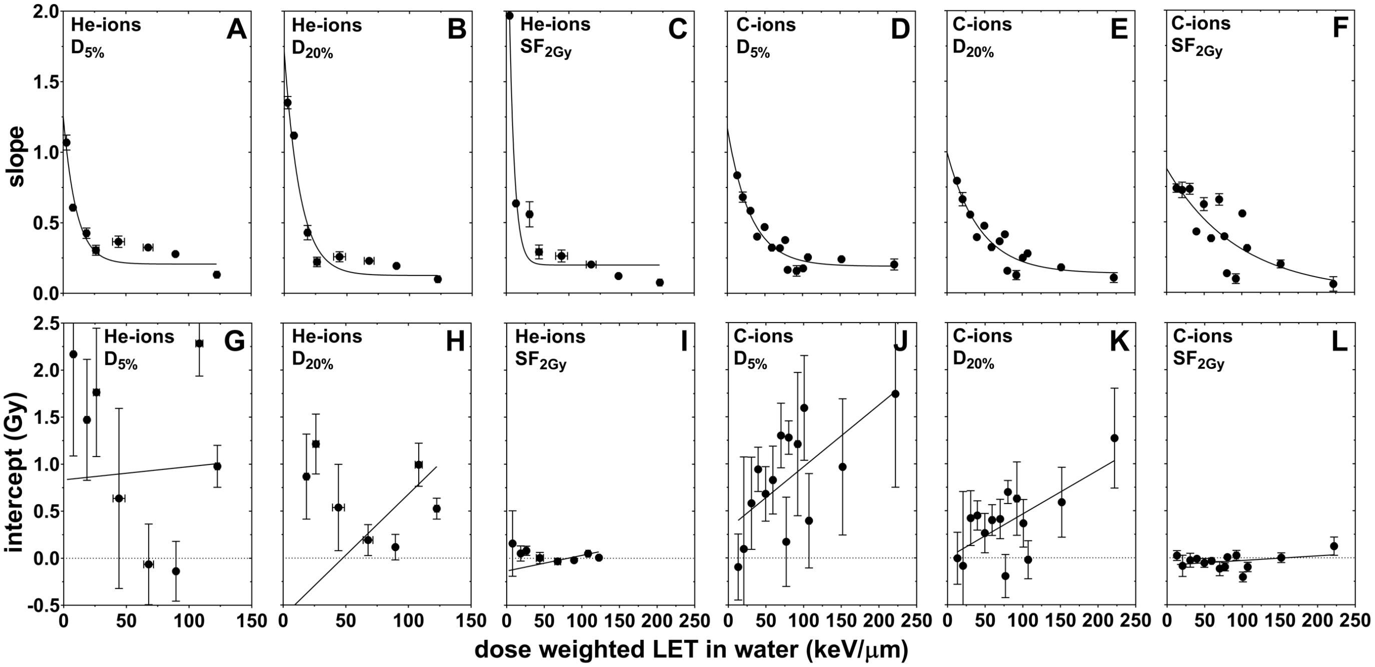


Supplemental Figure 3: Slope (A-F) and intercept (G-L) of the linear trends in the radiosensitivity parameters $D_{5\%}$ (A,G,D,J), $D_{20\%}$ (B,H,E,K),and $SF_{2Gy}$ (C,I,F,L) for the data obtained from the PIDE database (13) versus LET, binned by LET as described in Supplemental Method 6.

Supplemental Table 8 shows the goodness of fit parameters for the linear and exponential functions used to model the variation of the intercept and slope, respectively, as a function of LET:

Supplemental Table 8: $R^2$ values for the fits of the binned PIDE database data

| **Ion** | **Parameter** | **$R^2$ value** | | | | | |
|---|---|---|---|---|---|---|---|
| | | **$D_{5\%}$** | **$D_{10\%}$** | **$D_{20\%}$** | **$D_{37\%}$** | **$D_{50\%}$** | **$SF_{2Gy}$** |
| **He-ions** | *Slope* | 0.8411 | 0.8663 | 0.9576 | 0.9682 | 0.9733 | 0.9363 |
| | *Intercept* | 0.003186 | 0.0008205 | 0.2524 | 0.1037 | 0.09708 | 0.08689 |
| **C-ions** | *Slope* | 0.9182 | 0.8981 | 0.8768 | 0.8580 | 0.8329 | 0.6327 |
| | *Intercept* | 0.4198 | 0.4578 | 0.4622 | 0.4213 | 0.337 | 0.1205 |

**Supplemental Results 4: Model Parameters determined from the PIDE database**

Our formalism can thus be generalized into the following function of 5 parameters, c d f g and h, that predicts $R_{ion}$ from $R_{photon}$ for a given LET:

$$R_{ion} = \left[(c-d)\cdot e^{-f\cdot LET} - d\right]\cdot R_{photon} + g\cdot LET + h$$

With this form, there is no need to bin the data from the PIDE database to determine the values of c, d, f, g and h. Instead, we fit the measured response for all of the cell lines in the database for all LET values to this function.

Supplemental Table 9 and Supplemental Table 10 show the values determined for that fit while Supplemental Table 11 gives the $R^2$ values for these fits.

Supplemental Table 9: Model parameters fit to the He-ion data in the PIDE database.

| Parameter | $D_{5\%}$ | $D_{10\%}$ | $D_{20\%}$ | $D_{37\%}$ | $D_{50\%}$ | $SF_{2Gy}$ |
|---|---|---|---|---|---|---|
| c | 1.00635E+00 | 1.01931E+00 | 1.02079E+00 | 1.01449E+00 | 1.00679E+00 | 1.10072E+00 |
| $\sigma_c$ | 7.59806E-02 | 7.05671E-02 | 6.63710E-02 | 6.45857E-02 | 6.52721E-02 | 8.15166E-02 |
| d | 2.22046E-14 | 1.61043E-02 | 3.46806E-02 | 4.83522E-02 | 5.36863E-02 | 8.18826E-02 |
| $\sigma_d$ | 6.85953E-02 | 6.12812E-02 | 5.37049E-02 | 4.75243E-02 | 4.46277E-02 | 6.59951E-02 |
| f | 2.78642E-02 | 3.02113E-02 | 3.41113E-02 | 3.94856E-02 | 4.36078E-02 | 3.88516E-02 |
| $\sigma_f$ | 5.93737E-03 | 6.03079E-03 | 6.51352E-03 | 7.50243E-03 | 8.52589E-03 | 9.04455E-03 |
| g | 1.33601E-02 | 9.85123E-03 | 6.06519E-03 | 2.99009E-03 | 1.62235E-03 | 2.72189E-04 |
| $\sigma_g$ | 5.03605E-03 | 3.86446E-03 | 2.75224E-03 | 1.82310E-03 | 1.35787E-03 | 4.25193E-04 |
| h | 2.69752E-01 | 2.15190E-01 | 2.17120E-01 | 2.12625E-01 | 1.95766E-01 | 2.25285E-14 |
| $\sigma_h$ | 5.18587E-01 | 3.90362E-01 | 2.73556E-01 | 1.79175E-01 | 1.33004E-01 | 3.95650E-02 |
| $\sigma_{cd}$ | -1.64846E-04 | 5.22983E-05 | 2.40446E-04 | 3.43215E-04 | 3.80700E-04 | 4.35334E-04 |
| $\sigma_{cf}$ | -5.95678E-05 | -2.83690E-05 | 1.21112E-05 | 6.67079E-05 | 1.17452E-04 | 8.60909E-05 |
| $\sigma_{cg}$ | 1.97943E-04 | 1.26258E-04 | 7.04318E-05 | 3.48081E-05 | 2.08920E-05 | 1.04016E-05 |
| $\sigma_{ch}$ | -2.73269E-02 | -1.76511E-02 | -1.01639E-02 | -5.27511E-03 | -3.30850E-03 | -1.50685E-03 |
| $\sigma_{df}$ | 2.92456E-04 | 2.51946E-04 | 2.18859E-04 | 2.00369E-04 | 1.98350E-04 | 3.62244E-04 |
| $\sigma_{dg}$ | -2.61629E-04 | -1.75584E-04 | -1.05083E-04 | -5.82526E-05 | -3.91040E-05 | -2.01371E-05 |
| $\sigma_{dh}$ | 1.17826E-02 | 6.84343E-03 | 3.19652E-03 | 1.27075E-03 | 6.69713E-04 | 5.24683E-04 |
| $\sigma_{fg}$ | -2.35234E-05 | -1.81039E-05 | -1.36524E-05 | -1.01494E-05 | -8.41807E-06 | -2.90733E-06 |
| $\sigma_{fh}$ | 2.17196E-03 | 1.64063E-03 | 1.21876E-03 | 9.01811E-04 | 7.52211E-04 | 2.40274E-04 |
| $\sigma_{gh}$ | -2.16797E-03 | -1.23253E-03 | -6.03148E-04 | -2.57640E-04 | -1.41760E-04 | -1.31958E-05 |

Supplemental Table 10: Model parameters fit to the C-ion data in the PIDE database.

| Parameter | $D_{5\%}$ | $D_{10\%}$ | $D_{20\%}$ | $D_{37\%}$ | $D_{50\%}$ | $SF_{2Gy}$ |
|---|---|---|---|---|---|---|
| c | 9.81377E-01 | 9.74305E-01 | 9.52298E-01 | 9.17992E-01 | 8.87838E-01 | 9.51203E-01 |
| $\sigma_c$ | 3.05133E-02 | 3.02111E-02 | 3.16965E-02 | 3.55268E-02 | 3.92012E-02 | 5.38453E-02 |
| d | 1.27490E-01 | 1.32391E-01 | 1.36955E-01 | 1.27348E-01 | 1.10784E-01 | 1.16750E-01 |
| $\sigma_d$ | 5.21650E-02 | 5.16260E-02 | 5.27049E-02 | 5.79653E-02 | 6.41201E-02 | 9.62528E-02 |
| f | 1.88337E-02 | 1.88554E-02 | 1.91784E-02 | 1.91243E-02 | 1.87219E-02 | 1.89489E-02 |
| $\sigma_f$ | 2.25968E-03 | 2.29477E-03 | 2.53650E-03 | 2.93669E-03 | 3.25190E-03 | 4.40507E-03 |
| g | 7.44330E-03 | 5.34081E-03 | 3.21779E-03 | 1.79803E-03 | 1.25471E-03 | 2.22045E-14 |
| $\sigma_g$ | 2.32143E-03 | 1.91377E-03 | 1.53382E-03 | 1.19263E-03 | 9.89918E-04 | 3.39633E-04 |
| h | 6.24177E-02 | 2.27503E-14 | 2.22045E-14 | 2.22045E-14 | 2.22045E-14 | 2.22045E-14 |
| $\sigma_h$ | 1.84107E-01 | 1.50871E-01 | 1.20588E-01 | 9.35565E-02 | 7.75941E-02 | 2.64963E-02 |
| $\sigma_{cd}$ | -7.23998E-05 | -3.53523E-05 | 4.40604E-05 | 1.56368E-04 | 2.61799E-04 | 1.73632E-04 |
| $\sigma_{cf}$ | 4.42500E-06 | 6.89964E-06 | 1.31402E-05 | 2.44425E-05 | 3.55816E-05 | 3.74410E-05 |
| $\sigma_{cg}$ | 2.78286E-05 | 2.15972E-05 | 1.62294E-05 | 1.26206E-05 | 1.09591E-05 | 5.83268E-06 |
| $\sigma_{ch}$ | -3.52901E-03 | -2.76583E-03 | -2.16580E-03 | -1.74955E-03 | -1.53778E-03 | -7.96143E-04 |
| $\sigma_{df}$ | 1.05990E-04 | 1.06811E-04 | 1.20787E-04 | 1.54498E-04 | 1.90060E-04 | 3.83238E-04 |
| $\sigma_{dg}$ | -1.06754E-04 | -8.67180E-05 | -7.02442E-05 | -5.91076E-05 | -5.34206E-05 | -2.85935E-05 |
| $\sigma_{dh}$ | 5.15777E-03 | 4.19401E-03 | 3.38432E-03 | 2.86276E-03 | 2.60998E-03 | 1.34798E-03 |
| $\sigma_{fg}$ | -4.12412E-06 | -3.42170E-06 | -2.98629E-06 | -2.61233E-06 | -2.32981E-06 | -1.16364E-06 |
| $\sigma_{fh}$ | 2.61357E-04 | 2.14379E-04 | 1.84539E-04 | 1.58603E-04 | 1.39431E-04 | 7.16635E-05 |
| $\sigma_{gh}$ | -3.42584E-04 | -2.32121E-04 | -1.49011E-04 | -9.03178E-05 | -6.24330E-05 | -7.16324E-06 |

Supplemental Table 11: $R^2$ values for the fits of the PIDE database data without binning

| Ion | $R^2$ value | | | | | |
|---|---|---|---|---|---|---|
| | $D_{5\%}$ | $D_{10\%}$ | $D_{20\%}$ | $D_{37\%}$ | $D_{50\%}$ | $SF_{2Gy}$ |
| **He-ions** | 0.7422 | 0.7657 | 0.7820 | 0.7885 | 0.7853 | 0.7385 |
| **C-ions** | 0.8394 | 0.8435 | 0.8273 | 0.7928 | 0.7588 | 0.6068 |

Since our data agreed with the data in the PIDE database, we subsequently included our data into the database and recalculated the fits with higher statistical power. Those results are shown Supplemental Table 12 and Supplemental Table 13 while Supplemental Table 14 gives the $R^2$ values for these fits.

Supplemental Table 12: Model parameters fit to the He-ion data in the PIDE database combined with the data collected in this work.

| Parameter | $D_{5\%}$ | $D_{10\%}$ | $D_{20\%}$ | $D_{37\%}$ | $D_{50\%}$ | $SF_{2Gy}$ |
|---|---|---|---|---|---|---|
| c | 1.04780E+00 | 1.04072E+00 | 1.02178E+00 | 9.90923E-01 | 9.65001E-01 | 1.09046E+00 |
| $\sigma_c$ | 5.42169E-02 | 5.09764E-02 | 4.90772E-02 | 4.95474E-02 | 5.15459E-02 | 6.33957E-02 |
| d | 2.22045E-14 | 7.78489E-03 | 2.76345E-02 | 4.24203E-02 | 4.84719E-02 | 7.86959E-02 |
| $\sigma_d$ | 6.53046E-02 | 5.93334E-02 | 5.27288E-02 | 4.78505E-02 | 4.59050E-02 | 6.48099E-02 |
| f | 2.65441E-02 | 2.82650E-02 | 3.15239E-02 | 3.57642E-02 | 3.88637E-02 | 3.62921E-02 |
| $\sigma_f$ | 4.24151E-03 | 4.31749E-03 | 4.74185E-03 | 5.56270E-03 | 6.39743E-03 | 6.88036E-03 |
| g | 1.47517E-02 | 1.09784E-02 | 6.70673E-03 | 3.28864E-03 | 1.76961E-03 | 2.67036E-04 |
| $\sigma_g$ | 4.02525E-03 | 3.15552E-03 | 2.30860E-03 | 1.58750E-03 | 1.21604E-03 | 3.65781E-04 |
| h | 3.87311E-02 | 7.14691E-02 | 1.37130E-01 | 1.76450E-01 | 1.78329E-01 | 2.24264E-14 |
| $\sigma_h$ | 3.61239E-01 | 2.78172E-01 | 2.01132E-01 | 1.37450E-01 | 1.05327E-01 | 3.00307E-02 |
| $\sigma_{cd}$ | -2.02008E-05 | 5.40244E-05 | 1.52981E-04 | 2.23594E-04 | 2.58969E-04 | 3.11745E-04 |
| $\sigma_{cf}$ | -8.81988E-06 | 1.74189E-06 | 1.98449E-05 | 4.90732E-05 | 7.90990E-05 | 7.33837E-05 |
| $\sigma_{cg}$ | 1.02056E-04 | 6.97653E-05 | 4.30288E-05 | 2.50679E-05 | 1.75488E-05 | 7.08048E-06 |
| $\sigma_{ch}$ | -1.39647E-02 | -9.57731E-03 | -6.05681E-03 | -3.64044E-03 | -2.59697E-03 | -1.00487E-03 |
| $\sigma_{df}$ | 2.04431E-04 | 1.83395E-04 | 1.68592E-04 | 1.66333E-04 | 1.73543E-04 | 2.87964E-04 |
| $\sigma_{dg}$ | -2.09229E-04 | -1.47868E-04 | -9.37433E-05 | -5.64114E-05 | -4.02711E-05 | -1.82826E-05 |
| $\sigma_{dh}$ | 6.72822E-03 | 4.36331E-03 | 2.30969E-03 | 1.11899E-03 | 6.95149E-04 | 3.84475E-04 |
| $\sigma_{fg}$ | -1.23590E-05 | -9.76021E-06 | -7.69027E-06 | -6.01748E-06 | -5.16469E-06 | -1.76251E-06 |
| $\sigma_{fh}$ | 9.15949E-04 | 7.12767E-04 | 5.55472E-04 | 4.34717E-04 | 3.77101E-04 | 1.18887E-04 |
| $\sigma_{gh}$ | -1.08785E-03 | -6.50527E-04 | -3.39318E-04 | -1.57972E-04 | -9.26442E-05 | -7.87203E-06 |

Supplemental Table 13: Model parameters fit to the C-ion data in the PIDE database combined with the data collected in this work.

| Parameter | $D_{5\%}$ | $D_{10\%}$ | $D_{20\%}$ | $D_{37\%}$ | $D_{50\%}$ | $SF_{2Gy}$ |
|---|---|---|---|---|---|---|
| c | 9.79999E-01 | 9.81562E-01 | 9.74292E-01 | 9.55371E-01 | 9.31113E-01 | 1.00170E+00 |
| $\sigma_c$ | 2.71088E-02 | 2.70109E-02 | 2.88434E-02 | 3.33777E-02 | 3.80019E-02 | 5.05182E-02 |
| d | 1.30387E-01 | 1.32748E-01 | 1.32735E-01 | 1.24134E-01 | 1.18378E-01 | 1.13124E-01 |
| $\sigma_d$ | 5.13959E-02 | 5.13087E-02 | 5.36608E-02 | 5.95739E-02 | 6.39276E-02 | 9.93132E-02 |
| f | 1.87834E-02 | 1.86770E-02 | 1.88170E-02 | 1.90323E-02 | 1.94642E-02 | 1.86507E-02 |
| $\sigma_f$ | 2.15123E-03 | 2.14926E-03 | 2.33708E-03 | 2.75347E-03 | 3.22089E-03 | 4.00716E-03 |
| g | 6.83433E-03 | 5.03179E-03 | 3.25979E-03 | 1.84313E-03 | 1.04647E-03 | 2.22045E-14 |
| $\sigma_g$ | 2.25507E-03 | 1.87041E-03 | 1.52901E-03 | 1.21525E-03 | 1.01324E-03 | 3.44247E-04 |
| h | 1.38841E-01 | 3.99502E-02 | 2.70079E-14 | 2.22292E-14 | 2.59148E-02 | 2.22045E-14 |
| $\sigma_h$ | 1.71475E-01 | 1.41871E-01 | 1.16074E-01 | 9.26400E-02 | 7.77626E-02 | 2.59273E-02 |
| $\sigma_{cd}$ | -8.05359E-05 | -6.13477E-05 | -1.11549E-05 | 9.95318E-05 | 2.61539E-04 | 1.32756E-08 |
| $\sigma_{cf}$ | 2.40485E-06 | 3.85759E-06 | 7.96358E-06 | 1.79436E-05 | 3.26373E-05 | 2.32099E-05 |
| $\sigma_{cg}$ | 2.43720E-05 | 1.96556E-05 | 1.59711E-05 | 1.30701E-05 | 1.08440E-05 | 6.07599E-06 |
| $\sigma_{ch}$ | -2.99119E-03 | -2.41558E-03 | -2.01297E-03 | -1.72549E-03 | -1.52300E-03 | -7.81126E-04 |
| $\sigma_{df}$ | 1.00119E-04 | 1.00104E-04 | 1.14067E-04 | 1.49501E-04 | 1.87665E-04 | 3.61881E-04 |
| $\sigma_{dg}$ | -1.02586E-04 | -8.45741E-05 | -7.16056E-05 | -6.20522E-05 | -5.44336E-05 | -2.99997E-05 |
| $\sigma_{dh}$ | 4.74017E-03 | 3.93633E-03 | 3.34649E-03 | 2.91246E-03 | 2.54944E-03 | 1.36691E-03 |
| $\sigma_{fg}$ | -3.84567E-06 | -3.15652E-06 | -2.76151E-06 | -2.51204E-06 | -2.38077E-06 | -1.07676E-06 |
| $\sigma_{fh}$ | 2.32039E-04 | 1.89318E-04 | 1.64395E-04 | 1.48050E-04 | 1.39288E-04 | 6.35851E-05 |
| $\sigma_{gh}$ | -3.07186E-04 | -2.11691E-04 | -1.42104E-04 | -9.04873E-05 | -6.33960E-05 | -7.05699E-06 |

Supplemental Table 14: $R^2$ values for the fits of the unbinned PIDE database data combined with the data collected in this work.

| Ion | $R^2$ value | | | | | |
|---|---|---|---|---|---|---|
| | $D_{5\%}$ | $D_{10\%}$ | $D_{20\%}$ | $D_{37\%}$ | $D_{50\%}$ | $SF_{2Gy}$ |
| **He-ions** | 0.8374 | 0.8505 | 0.8556 | 0.8484 | 0.8349 | 0.8022 |
| **C-ions** | 0.8542 | 0.8597 | 0.8453 | 0.8069 | 0.7640 | 0.6297 |

As a general note, after incorporating our data into the training data, the uncertainties in the model parameters are lower, and the $R^2$ values of the fits are higher, implying an improved performance of the model after incorporating our additional data into the training data.

**Supplemental Results 5: $R_{ion}$ versus $R_{photon}$ predictions**

Using our model, we predicted the relationship between ion and photon radiosensitivity for 2.2, 7.0 and 14.0 keV/µm for He-ions and 13.5, 27.9 keV/µm and 60.5 keV/µm for C-ions, for the parameters $D_{10\%}$, $D_{37\%}$ and $D_{50\%}$ (Fig 3). The predictions for the parameters $D_{5\%}$, $D_{20\%}$ and $SF_{2Gy}$ are given in Supplemental Figure 4.

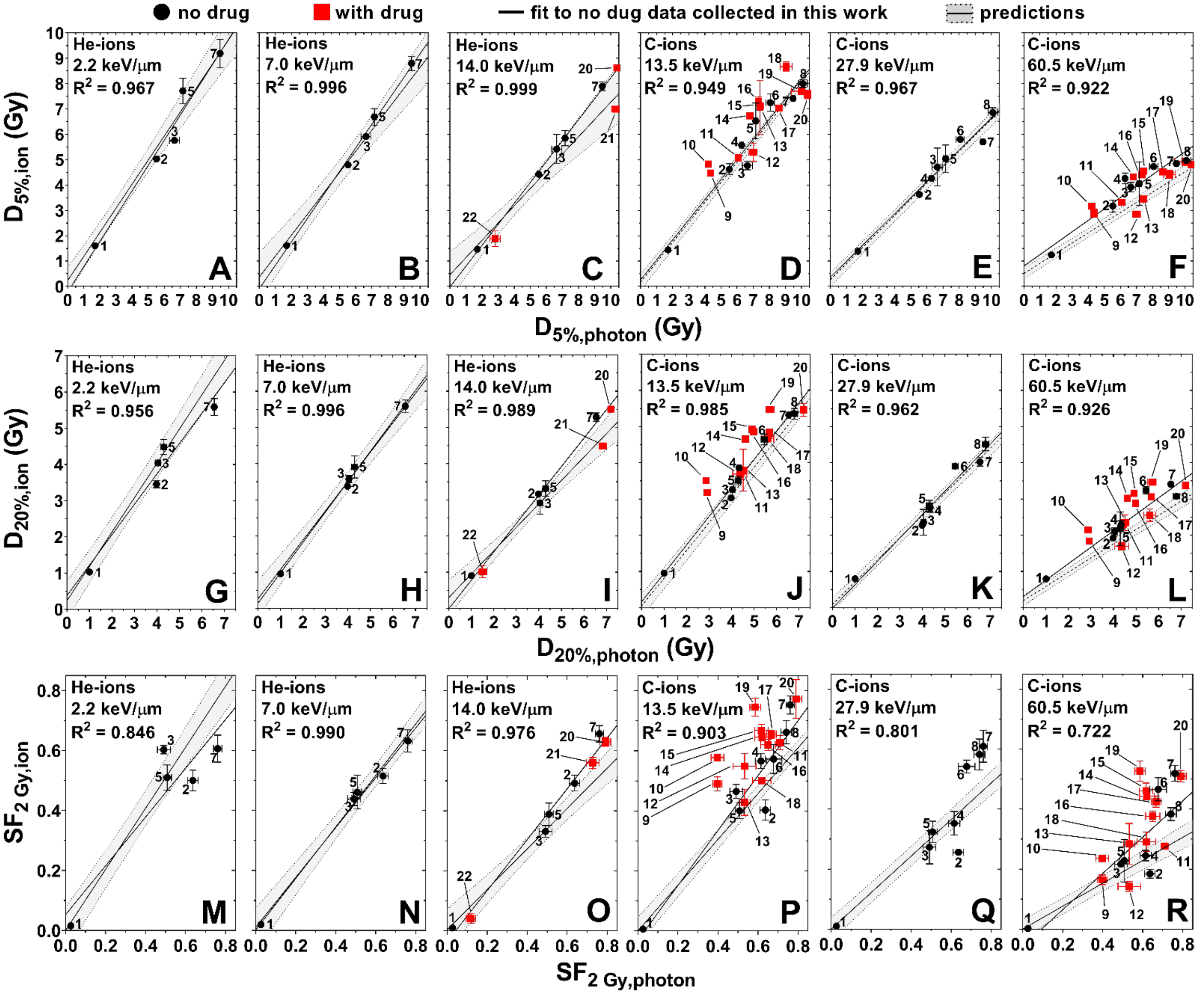


Supplemental Figure 4: Prediction (with 95% confidence interval) and measured data for the linear relation between $D_{5\%}$ (A-F), $D_{20\%}$ (G-L) and $SF_{2Gy}$ (M-R) for ions vs. photons. Numbers indicate the cell line and are given in caption of Fig. 1.

Using our model, we were able to predict the $R_{ion}$ versus $R_{photon}$ trend for three He-ion LET values (2.2, 7.0 and 14.0 keV/µm) and three C-ion LET values (13.5, 27.9 and 60.5 keV/µm). Supplemental Table 15 summarizes the accuracy of those predictions.

Supplemental Table 15: Root mean square percentage error between predicted $R_{ion}$ and $R_{photon}$ trend and values derived from survival data for cells exposed to He-, C-ions and 6 MV x-rays

| Root mean square percentage error | | | | | | | |
|---|---|---|---|---|---|---|---|
| LET / Parameter | He-ions | | | C-ions | | | Total Combined |
| | 2.2 keV/µm | 7.0 keV/µm | 14.0 keV/µm | 13.5 keV/µm | 27.9 keV/µm | 60.5 keV/µm | |
| $D_{5\%}$ | 10.7% | 6.1% | 9.9% | 8.6% | 5.3% | 21.8% | 12.3% |
| $D_{10\%}$ | 9.6% | 5.1% | 8.9% | 7.3% | 6.0% | 21.3% | 11.7% |
| $D_{20\%}$ | 10.7% | 5.6% | 9.5% | 9.7% | 8.4% | 23.4% | 13.3% |
| $D_{37\%}$ | 15.1% | 7.9% | 11.7% | 17.3% | 11.7% | 29.6% | 18.0% |
| $D_{50\%}$ | 19.8% | 10.1% | 13.8% | 24.8% | 16.4% | 35.2% | 22.9% |
| $SF_{2Gy}$ | 25.0% | 9.4% | 28.4% | 31.1% | 26.4% | 51.1% | 32.7% |
| Total Combined | 16.2% | 7.6% | 15.3% | 18.7% | 14.3% | 32.1% | |

**Supplemental Results 6: Accuracy of RBE versus $R_{photon}$ predictions**

Equation 3 predicts the RBE for a given ion LET for a particular cell line based upon its radiosensitivity. The predictions for $RBE_{D10\%}$, $RBE_{D37\%}$ and $RBE_{D50\%}$ are given in Fig. 4, while the predictions for $RBE_{D5\%}$ and $RBE_{D20\%}$ are given in Supplemental Figure 5.

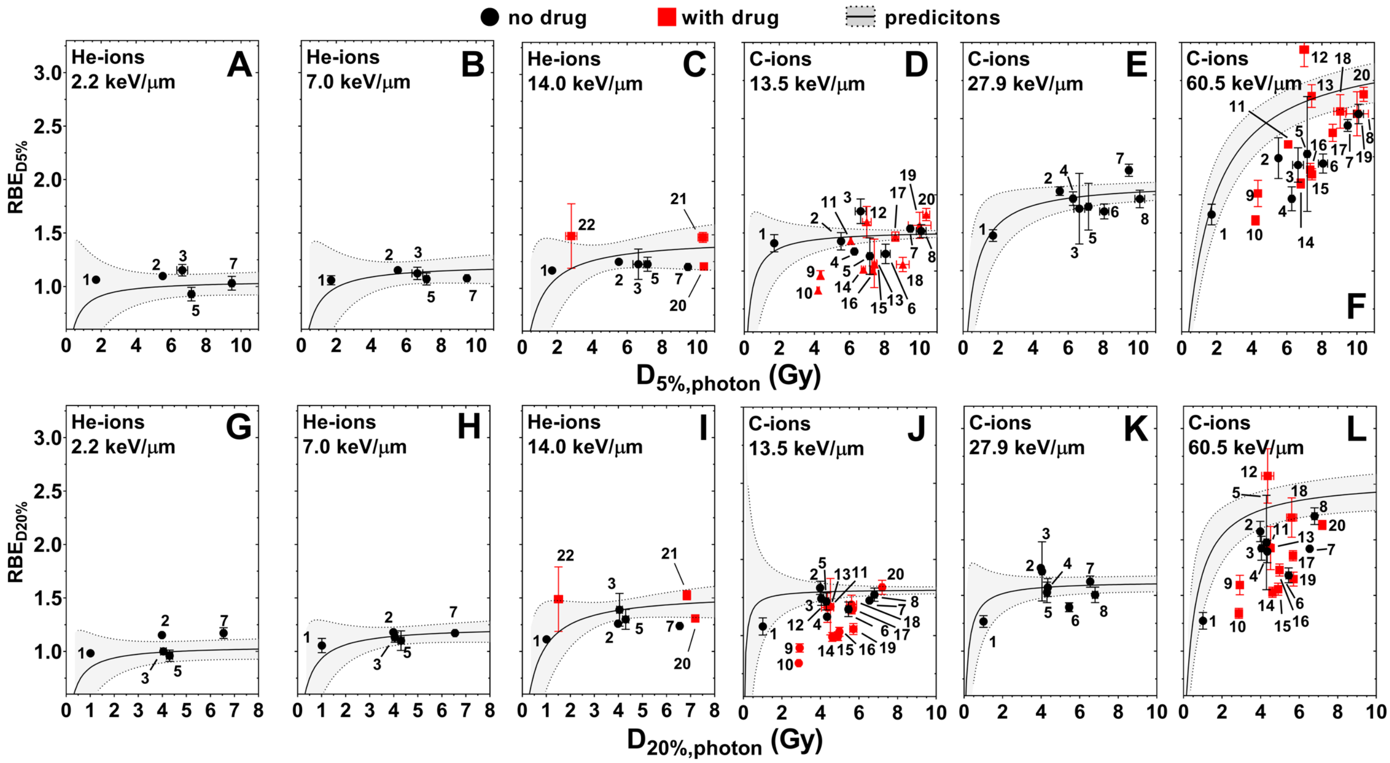


Supplemental Figure 5: Predicted (with 95% confidence interval) and measured RBE as functions of $D_{5\%,photon}$ (A-F) and $D_{20\%,photon}$ (G-L). Numbers indicate the cell line and are given in the caption of Fig. 1. These cell lines and LET values were not used to determine the linear functions and served as a validation of the model. The RBE values predicted for $D_{10\%}$, $D_{37\%}$ and $D_{50\%}$ are in given in Fig. 4.

Using our model, we were able to predict the RBE versus $R_{photon}$ trend for three He-ion LET values (2.2, 7.0 and 14.0 keV/μm) and three C-ion LET values (13.5, 27.9 and 60.5 keV/μm). Supplemental Table 16 summarizes the accuracy of those predictions.

Supplemental Table 16: Root mean square percentage error between predicted RBE and $R_{photon}$ trend and values derived from survival data for cells exposed to He-, C-ions and 6 MV x-rays.

| Root mean square percentage error | | | | | | | |
|---|---|---|---|---|---|---|---|
| LET | He-ions | | | C-ions | | | Total Combined |
| Parameter | 2.2 keV/μm | 7.0 keV/μm | 14.0 keV/μm | 13.5 keV/μm | 27.9 keV/μm | 60.5 keV/μm | |
| RBE ($D_{5\%}$) | 8.1% | 6.2% | 9.3% | 8.4% | 5.4% | 17.0% | 10.2% |
| RBE ($D_{10\%}$) | 5.0% | 5.3% | 8.1% | 6.8% | 5.7% | 17.1% | 9.5% |
| RBE ($D_{20\%}$) | 6.6% | 6.2% | 8.5% | 8.5% | 8.1% | 18.1% | 10.7% |
| RBE ($D_{37\%}$) | 13.4% | 9.2% | 10.3% | 13.9% | 11.6% | 20.5% | 14.2% |
| RBE ($D_{50\%}$) | 18.8% | 12.4% | 12.0% | 18.5% | 17.9% | 22.9% | 18.1% |
| Total Combined | 11.6% | 13.6% | 9.7% | 12.0% | 10.8% | 19.3% | / |

**Supplemental Results 7: Survival curves predicted for PANC-1, AsPC-1 and Panc 10.05 cell lines exposed to C-ion radiation**

As we did not expose the AsPC-1, PANC-1 and Panc 10.05 cell lines to He-ion radiation, but did expose them to C-ion radiation, we did not include their predicted survival curves in Fig. 5. Further, as we did include the survival curve predictions for the cells treated with DNA-repair inhibitors in Fig. 5 for brevity, these predictions are given in the supplemental figures below:

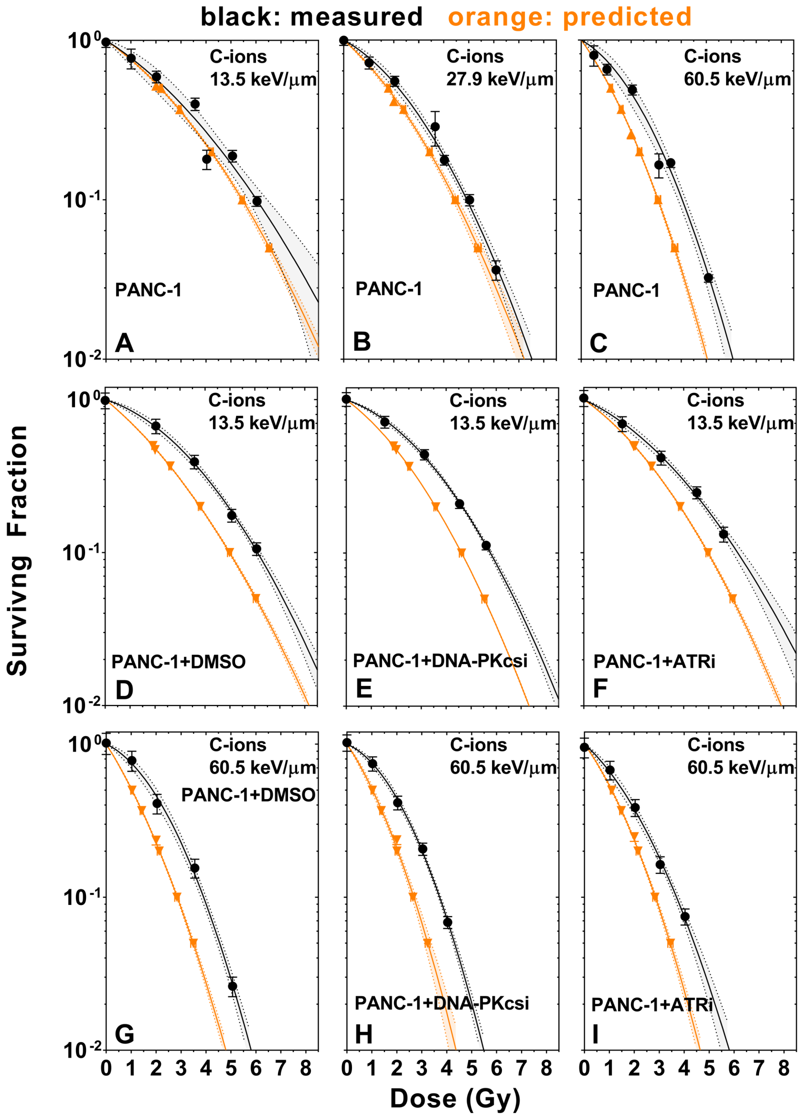


Supplemental Figure 6: Predicted (orange, with 95% confidence interval) and measured (black, with 95% confidence interval) survival curve for Panc1 cells exposed to C-ions with LETs ranging from 13.5 – 60.5 keV/µm. Cells were treated with the DNA repair inhibitors to inhibit DNA-PKcs (E,H), ATR (F,I), with just the DMSO vehicle (D,G), and without any treatment (A-C).

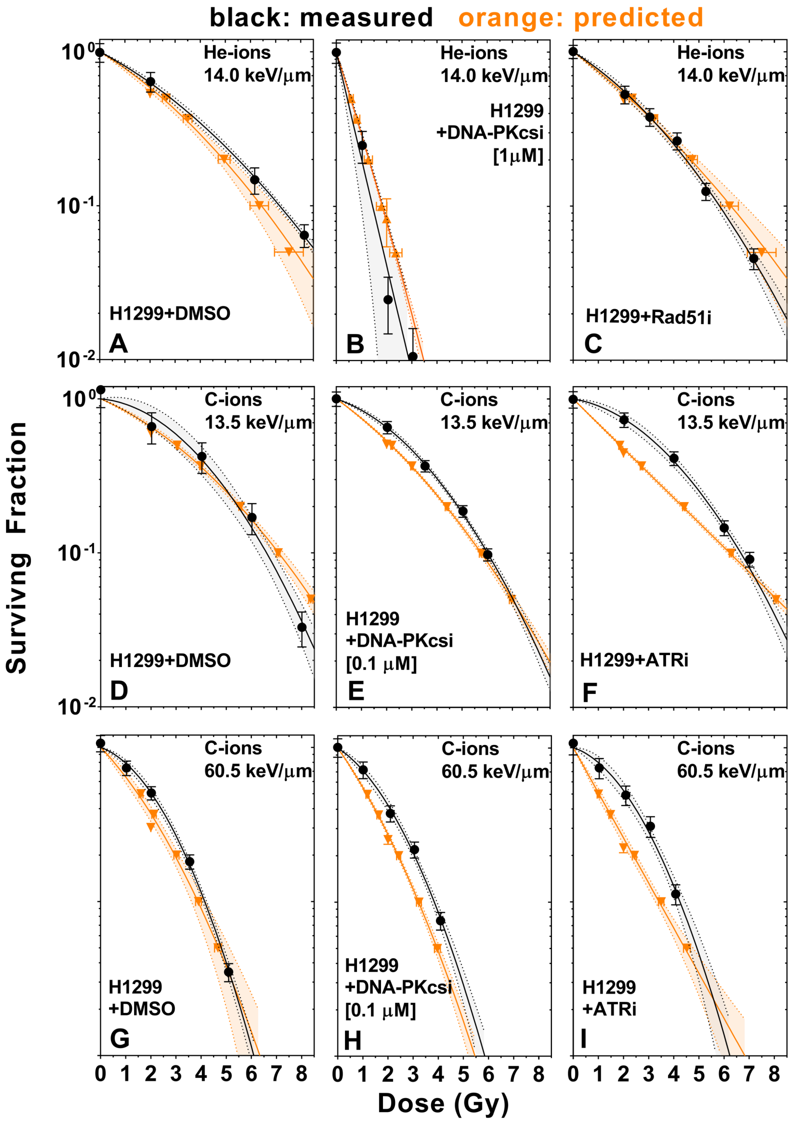


Supplemental Figure 7: Predicted (orange, with 95% confidence interval) and measured (black, with 95% confidence interval) survival curve for H1299 cells exposed to He- and C-ions with LETs ranging from 13.5 – 60.5 keV/µm. Cells were treated with the DNA repair inhibitors to inhibit DNA-PKcs (B,E,H), ATR (F,I), Rad51 (C) and with just the DMSO vehicle (A,D,G).

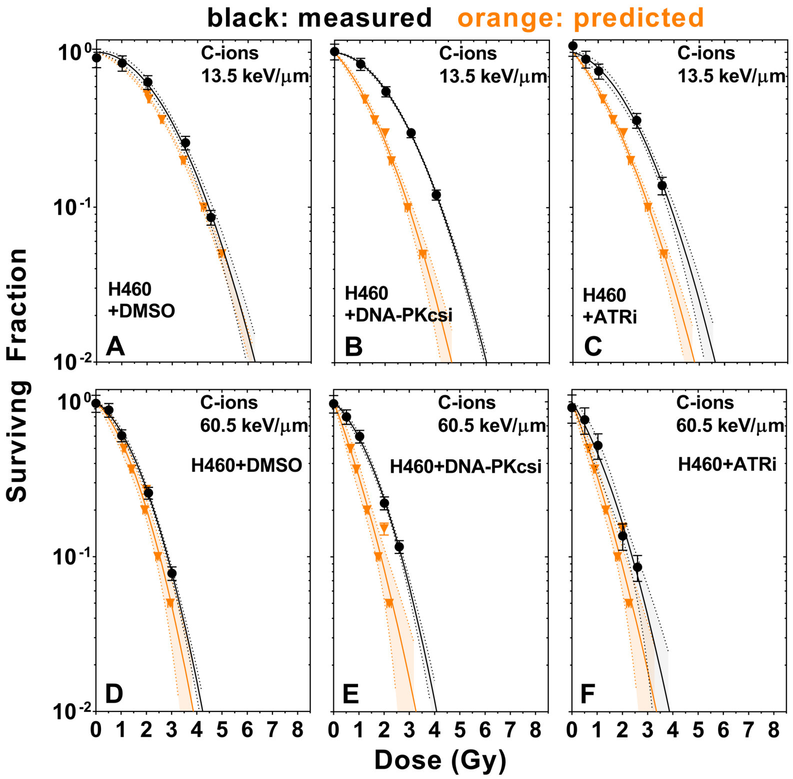


Supplemental Figure 8: Predicted (orange, with 95% confidence interval) and measured (black, with 95% confidence interval) survival curve for H460 cells exposed to C-ions with LETs of 13.5 and 60.5 keV/µm. Cells were treated with the DNA repair inhibitors to inhibit DNA-PKcs (B,E), ATR (C,F), and with just the DMSO vehicle (A,D).

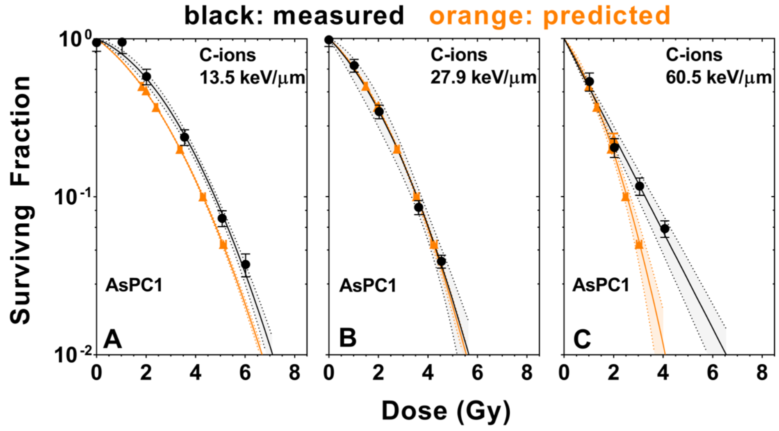


Supplemental Figure 9: Predicted (orange, with 95% confidence interval) and measured (black, with 95% confidence interval) survival curve for AsPC1 cells exposed to C-ions with LETs ranging from 13.5 – 60.5 keV/µm.

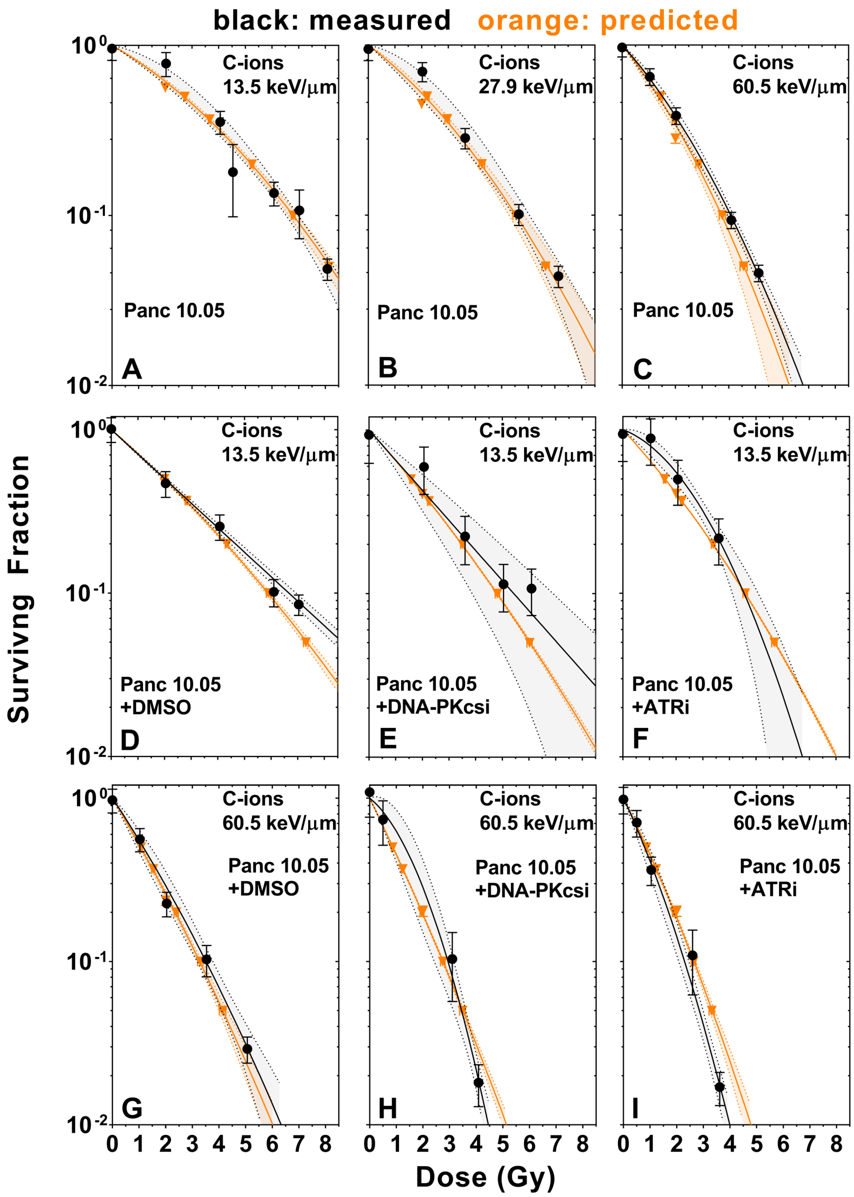


Supplemental Figure 10: Predicted (orange, with 95% confidence interval) and measured (black, with 95% confidence interval) survival curve for Panc105 cells exposed to C-ions with LETs ranging from 13.5 – 60.5 keV/µm. Cells were treated with the DNA repair inhibitors to inhibit DNA-PKcs (E,H), ATR (F,I), with just the DMSO vehicle (D,G), and without any treatment (A-C).

**Supplemental Results 8: Prediction of cell lines treated with ion and DNA repair inhibitors**

Supplemental Figure 11 shows the response of DNA repair-inhibited cells compared to the 95% prediction interval based upon the untreated cells for the radiosensitivity parameters $D_{5\%}$, $D_{20\%}$ and $SF_{2Gy}$. The data for the parameters $D_{10\%}$, $D_{37\%}$ and $D_{50\%}$ is given in Fig. 6.

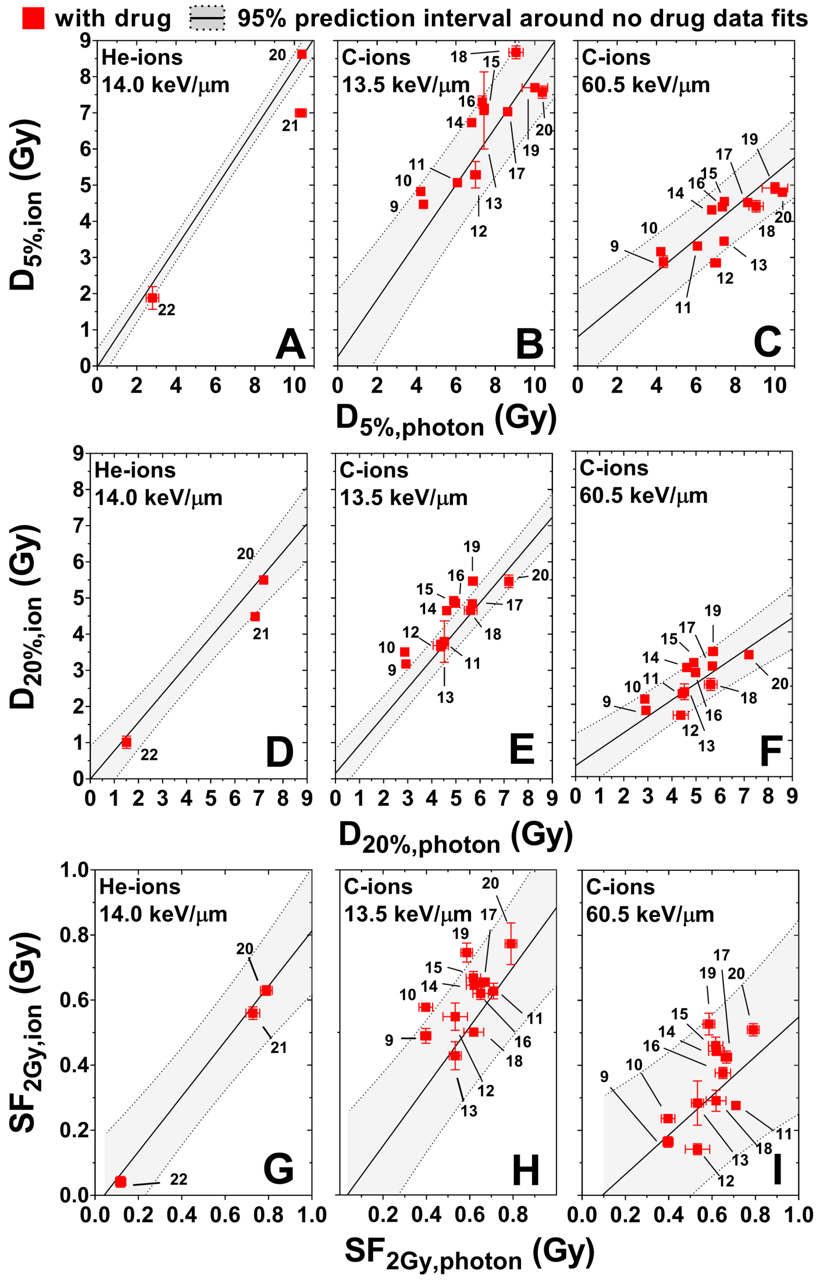


Supplemental Figure 11: Measured ion versus photon radiosensitivity for cells treated with DNA repair inhibitors (red) superimposed on the 95% prediction interval derived from the data not treated with drugs (gray) for the parameters $D_{5\%}$ (A-C), $D_{20\%}$ (D-F) and $SF_{2Gy}$ (G-I). Numbers that indicate the cell line and are given in the caption of Fig. 1. Results for $D_{10\%}$, $D_{37\%}$ and $D_{50\%}$ are given in Fig. 6.
